# Synaptic Targets of Circadian Clock Neurons Influence Core Clock Parameters

**DOI:** 10.1101/2025.01.30.635801

**Authors:** Eva Scholz-Carlson, Aishwarya Ramakrishnan Iyer, Aljoscha Nern, John Ewer, Maria P. Fernandez

## Abstract

Neuronal connectivity in the circadian clock network is essential for robust endogenous timekeeping. In the *Drosophila* circadian clock network, four pairs of small ventral lateral neurons (sLN_v_s) serve as critical pacemakers. Peptidergic communication via sLN_v_ release of the key output neuropeptide *Pigment Dispersing Factor* (PDF) has been well characterized. In contrast, little is known about the role of the synaptic connections that sLN_v_s form with downstream neurons. Connectomic analyses revealed that the sLN_v_s form strong synaptic connections with a group of previously uncharacterized neurons, SLP316. Here, we show that silencing synaptic output in the SLP316 neurons via tetanus toxin (TNT) expression shortens the free-running period, whereas hyper-exciting them by expressing the Na[+] channel NaChBac results in period lengthening. Under light-dark cycles, silencing SLP316 neurons also causes lower daytime activity and higher daytime sleep. Our results revealed that the main postsynaptic partners of the *Drosophila* pacemaker neurons are a non-clock neuronal cell type that regulates the timing of sleep and activity.

## Introduction

Circadian rhythms, which allow animals to adapt to day-night changes in their environment, are generated by small groups of neurons in the brain that contain molecular clocks (1). These neurons form a network with synchronized molecular oscillations (2, 3). The maintenance of robust and coherent behavioral circadian rhythms depends on connectivity within the clock network, which controls the timing of key behavioral outputs like rhythms in feeding, mating, and sleep (3). In both mammals and flies, connectivity in the clock network is mediated by peptide and transmitter outputs (4–7). Although the molecular mechanisms driving endogenous oscillations in clock neurons are well described, the pathways through which these neurons communicate with each other and with downstream targets to propagate oscillations throughout the brain remain poorly understood.

*Drosophila melanogaster* is an ideal model for studying connectivity in the clock network. The fly circadian network consists of ∼240 neurons ((8), reviewed in (9)) and is the functional equivalent of the mammalian suprachiasmatic nuclei, the principal clock in the vertebrate brain (10, 11). Each clock neuron within this network has an intracellular molecular timekeeping mechanism based on a transcriptional‒translational feedback loop (12). Clock neurons can be organized into seven main groups based on their anatomical location (11, 13–16). Whereas the molecular oscillations of most of these groups are in phase, some major clock groups become active at specific times of day, which correlates with their roles in behavioral outputs (17). Among these cells, the small lateral ventral neurons (sLN_v_s) are key for maintaining rhythmicity under conditions of constant darkness and temperature (DD) and are considered the dominant circadian pacemakers under these conditions (18–21). Despite the importance of these cells for sustaining physiological and behavioral rhythms, very little is known about their connectivity with target neurons. Indeed, an analysis of their synaptic connectivity revealed that they form very few monosynaptic connections with other clock neurons (16).

Peptidergic signaling is essential for clock network intercommunication and output (6). The neuropeptide *Pigment Dispersing Factor* (PDF), which is secreted by the sLN_v_s, is required for synchronizing molecular oscillations within the network, setting the phase of neural activity in other groups of clock neurons, and maintaining activity-rest rhythms under constant conditions (18, 22–24). Expression of PDF undergoes daily and circadian oscillations in the sLN_v_ dorsal terminals (19), and the loss of PDF or of the PDF receptor (PDFR) results in arrhythmicity under free-running conditions (25–27). In addition to PDF, the sLN_v_s express the neuropeptide, short Neuropeptide F (sNPF) (28), which promotes sleep and is involved in feeding regulation (28–30), and the inhibitory neurotransmitter glycine (31). Interestingly, downregulation of *bruchpilot* (*brp*) and of other synaptic active zone proteins expressed by the sLN_v_s does not affect rhythmicity, and dense-core vesicle release profiles of the sLN_v_s in the Full Adult Fly Brain (FAFB) do not show spatial overlap with BRP-labeled active zones (32). This finding is consistent with the observation that neuropeptide release takes place from sites located all along the sLN_v_, including the soma (33), and with the persistence of robust behavioral rhythmicity upon silencing of synaptic transmission (34) in *Pdf*+ neurons (35, 36)

Although the circadian clock provides predictability, it must also be flexible to adapt to varying environmental conditions. Distinct subsets of neurons control the morning and evening bouts of activity that produce the bimodal pattern that flies show under light-dark cycles. The sLN_v_s, which are active around dawn (17), are associated with the morning peak of activity (18, 20, 21) and are called morning (M) cells. The PDFR-expressing lateral neurons dorsal (LN_d_s) and the PDF-negative 5^th^ LN_v_ are associated with the evening peak activity (20, 21, 37) and their neural activity peaks before dusk (17). Some dorsal neurons (DNs) also contribute to the timing and amount of activity and sleep via the modulation of M and E cells (38–41). The phase of the morning and evening activity bouts adjusts to environmental light and temperature (42–47), allowing flies to adjust their activity patterns according to the season.

The four sLN_v_s form few direct synaptic connections with other clock neurons or with other neurons outside the clock network, yet they form strong connections with a previously uncharacterized group of neurons named SLP316. The SLP316 cells synapse onto the DN1pA class of dorsal clock neurons, which are associated with the reorganization of sleep/wake cycles (reviewed in (40)). Here, we show that despite not being clock neurons themselves, silencing SLP316 neuronal activity resulted in a shortening of the free-running period of activity, whereas their excitation resulted in the opposite phenotype. Neither manipulation led to the loss of rhythmicity. Under light-dark cycles (LD), manipulations of the SLP316 activity led to a reduction in evening activity and an increase in sleep, both under ambient temperature of 25°C, and under cold, winter-like days. Under both LD and DD, the behavioral phenotypes observed were different from those caused by the loss of PDF. In conclusion, our data reveal a role for a previously unknown neuronal population that connects two classes of clock neurons but is not itself part of the circadian clock network, in regulating the timing of sleep/wake cycles.

## Results

### 1 Connectomics datasets identify SLP316 neurons as primary post-synaptic partners of sLN_v_s

Based on the *Drosophila* hemibrain connectome (48), the sLN_v_s form much stronger connections with an uncharacterized neuronal cell type than with any other neurons, including other clock neurons (16) (Fig. 1A). This cell type, SLP316, consists of three neurons with somas in the dorsal-most region of the Superior Lateral Protocerebrum (SLP) (48). Interestingly, the sLN_v_s are in turn the main presynaptic inputs to SLP316 neurons (Fig 1A, B). Most inputs to SLP316 cells from their top ten presynaptic partners are located along the ventromedial projections, whereas their output synapses are concentrated along the dorsal arborizations (Fig. 1C).

**Fig. 1:**
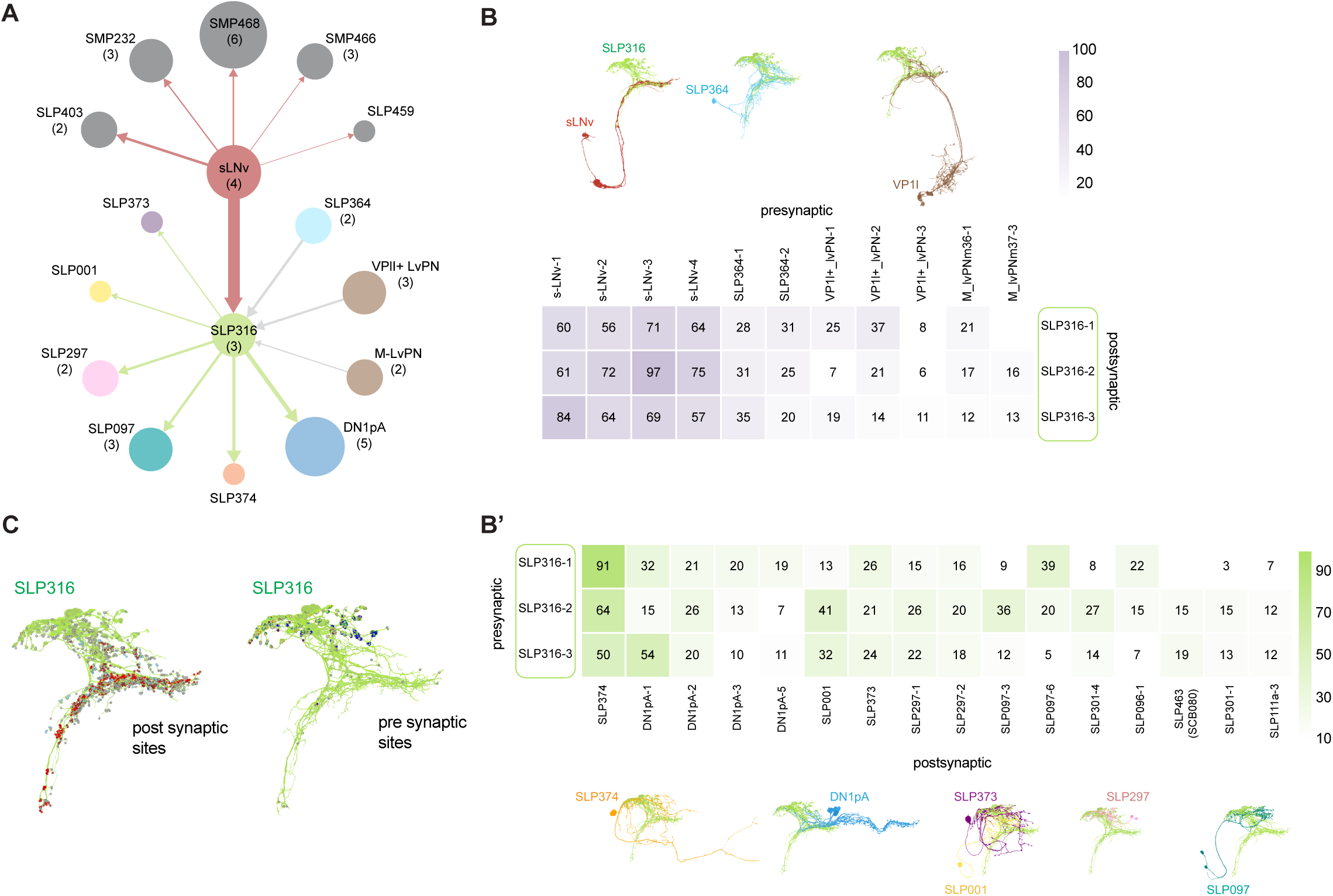
SLP316 neurons are the main postsynaptic partner of the sLN_v_s. (A-B) Connectomic mapping from the hemibrain dataset. (A) Map of top input/output neuronal groups to/from the SLP316s and from the sLN_v_s. The number of neurons in each group are indicated in parenthesis if greater than one and reflected by circle size. Connection strength between cell types is represented by arrow width and decreases in a clockwise direction for each connection type (input/output). SLP232 likely corresponds to DN3s. (B) Strong shared inputs to, and (B’) outputs of the SLP316s (>10 synapses to/from at least two SLP316 neurons) with neuronal morphology shown above or below. (C) Input synapses to, and output synapses of, the three SLP316 neurons (SLP316_R_1, SLP316_R_2 and SLP316_R_3) shown in green, excluding synapses from neuron segments. Synapse color corresponds to neuron color in (A) and (B) and all other synapses are shown in grey.

The synapses formed by the sLN_v_s onto the SLP316 neurons represent 70% of the synapses by all groups that form strong connections with these cells (weight >10) (Fig 1A, B). Many of the synapses that the sLN_v_s form onto the SLP316s (∼20%) are located below the point at which the sLN_v_s extend and ramify their dorsal termini (Suppl. Fig. 1A), the region long thought to be the location of sLN_v_ output synapses. The other main inputs to SLP316 are from two neurons that belong to the SLP364 cell type and from ventroposterior (VP) antennal lobe glomeruli neurons. The SLP316s are innervated largely by neurons in the SLP neuropil but also from the superior medial (SMP) and posterior lateral (SLP) protocerebrum (Suppl. Fig. 1B).

The SLP316 neurons are largely homogeneous in their connectivity patterns with upstream neurons, but their outputs are more varied (Fig 1B’). These cells form some of the strongest connections with another group of clock neurons, posterior dorsal neurons (DN1p). The DN1pA class of clock neurons is the top output of the SLP316 cells. The main neuropil innervated by SLP316s is the SLP, followed by the SMP and PLP with a much smaller number of connections (Suppl. Fig. 1B). We next examined the connectivity patterns of the sLN_v_s in the FlyWire *Drosophila* connectome (49) to determine whether cells similar in morphology to the SLP316s were among their top post-synaptic partners. We found that in both hemispheres, the sLN_v_s form most of their output synapses with cells highly similar in location and morphology (Suppl. Fig. 1C). Computational predictions based on synapse architecture for both the hemibrain and FAFB-Flywire datasets (50) indicate that SLP316 cells are likely glutamatergic.

### 2 The SLP316 neurons are non-clock neurons that extend along the sLN_v_ projection tract

To further study the SLP316 neurons, we generated a Split-Gal4 line that targets these cells (#SS76489, see Methods) (Suppl. Fig. 2A). Since both the hemibrain and FAFB/FlyWire connectomes correspond to an adult female (48), we dissected brains from both *SLP316>mCD8-GFP* males and females to determine if there were sex differences in the number of cells or their morphology. The projections from the SLP316 neurons extend toward the ventral brain along the same tract as the projections of the sLN_v_s, which extend from the ventral to the dorsal brain (Fig. 2A-B). We did not find expression in additional neurons in the central brain or the ventral nerve cord, or sex differences in the number of cells (Fig. 2C). The number of cells remains similar in older flies (Suppl. Fig. 2B). Therefore, we conclude that the driver is likely specifically expressed in SLP316 neurons in adults and that the expression pattern is similar in males and females.

**Fig. 2:**
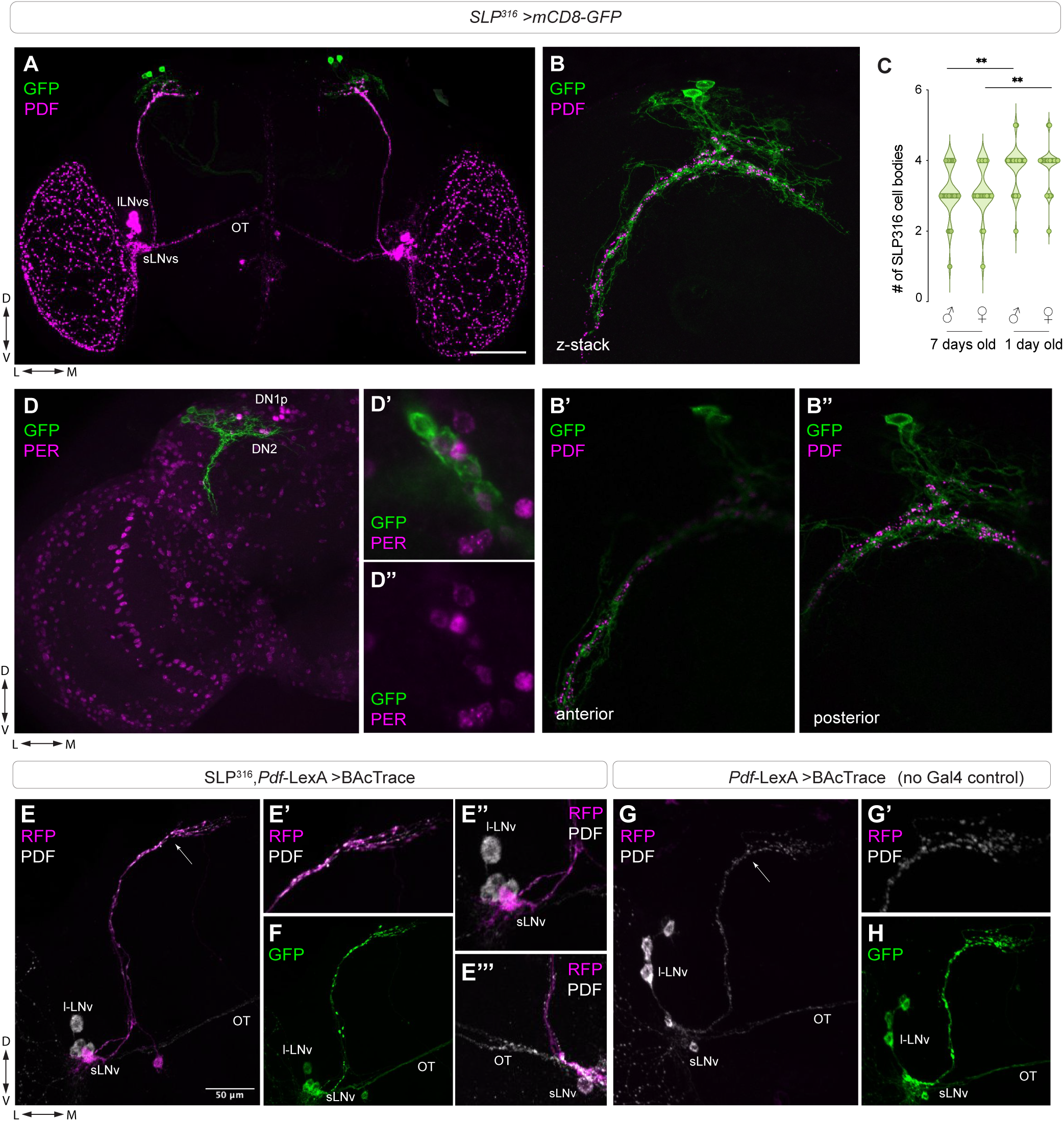
The SLP316 driver labels non-clock neurons that share functional synapses with the sLN_v_s. (A-B) Representative confocal images of a brain from a 7-day old male fly (*SLP316>GFP)* stained with antibodies against GFP (green) and PDF (magenta). (A) Whole brain, (B) Superior Lateral Protocerebrum z-stack, and anterior (B’) and posterior (B”) single optical slices (step size 0.9 μm). (C) Quantification of cell body count on the left hemisphere labelled by the *SLP316* driver in 7-day and 1-day old male and female flies from three replicate experiments (n > 21 for each age/sex). (D) Representative confocal images of a brain hemisphere of 6–7-day old male brains (*SLP316>GFP*) stained with antibodies against GFP (green) and PER (magenta). (E-F) Representative confocal images of *SLP316-Gal4* and *Pdf-LexA* driving expression of BAcTrace components. RFP signal can only be detected in the sLNvs. (E’) sLN_v_ dorsal termini show PDF and RFP expression. (E’’) Cell bodies of sLN_v_s and lLN_v_s. (E’’’) sLN_v_ cell bodies plus dorsal projections show RFP signal, whereas the lLN_v_s do not. (F) syb-GFP expression in the small and large LN_v_s in experimental brains. (G-H) Representative confocal images of *Pdf-LexA* driving expression of BATrace components, without the SLP316-Gal4 driver. All scale bars are 50 µm.

The SLP316 neurons are in the dorsoposterior area of the brain, where the DN1ps are located. To determine if these cells were yet unidentified clock neurons, we co-stained with a PER antibody. Adult *SLP316>mCD8-GFP* flies were entrained to a 12h:12h light-dark cycle (LD) for 5 days, and brains were dissected at the end of the night (ZT23), when PER nuclear signal is strongest (51). We did not detect PER nuclear signal in the somas of any of the SLP316 cells (Fig. 2D). Therefore, we conclude that the SLP316 neurons are not circadian clock neurons. To determine if there were daily changes in excitability in these cells, we used a transcriptional reporter in which the *Hr38* regulatory DNA is upstream of a nuclear-localized fluorescent Tomato reporter (52). In the sLN_v_s, the *Hr38-Tomato* reporter gene shows higher expression around ZT2, consistent with a previous study (53). In contrast, SLP316 cells showed consistently low signal, and the presence of physiological rhythms cannot be excluded based on these data (Suppl. Fig. 2C-D).

Next, we assessed connectivity between the cells labeled by the SLP316 split-GAL4 line and the sLN_v_s via BAcTrace. This is a retrograde synaptic tracing technique in which *Clostridium botulinum* neurotoxin A1 (BoNT/A) is transferred from a postsynaptic donor cell (driven by Gal4 components) to a presynaptic receiver cell (driven by LexA components) by targeting the interior of a synaptic vesicle (54). In the receiver cell, the toxin releases a membrane-localized transcription factor, inducing Tomato expression. When a vesicle is recycled, the toxin is internalized into the cell. We combined flies expressing the BAcTrace transgenes (see the methods section) with flies carrying both the SLP316 split Gal4 and the *Pdf-LexA* transgenes and stained brains with antibodies against PDF, RFP, and GFP (Fig 2E-H). Experimental flies showed RFP signal in the sLN_v_s but not the lLN_v_s, whereas PDF-immunoreactivity was detected in both groups of LN_v_s (Fig. 2E). Control flies without the SLP316 split Gal4 driver did not express RFP (Fig. 2G). As expected, GFP signal could be detected in both the sLN_v_s and lLN_v_s in control flies, reflecting the expression pattern of the *Pdf-LexA* driver present in both lines (Fig. 2F and H). We did not observe sex differences in expression pattern in experimental or control flies. These results support the notion that the SLP316 neurons are post-synaptic partners of the sLN_v_s, which is consistent with connectomics data.

### 3 Altering the activity of SLP316 neurons changes the free-running period

Elimination of the LN_v_s via expression of the pro-apoptotic gene *hid* leads to severe loss of rhythmicity and shortening of the free running period, similar to the behavioral phenotypes of the *Pdf* null mutation, *Pdf*^01^(18). Whereas inhibiting synaptic transmission in *Pdf* neurons via tetanus toxin (TNT) does not affect the ability of the flies to maintain rhythmicity in DD, it leads to changes in free-running period, as *Pdf*>TNT flies show longer periods than their parental controls (35, 36). Since the SLP316s are the main cell type downstream of the clock pacemaker neurons, we asked how manipulations of these cells would affect behavioral rhythms.

The bacterial voltage-gated sodium channel *NaChBac* is used to increase neuronal excitability (reviewed in (55)). Expression of *NaChBac* in *Pdf*+ expressing neurons results in complex rhythms, with multiple superimposed period components under free-running conditions (56). *NaChBac* expression in the SLP316 cells resulted in lengthening of the free-running period without affecting overall rhythmicity (Fig. 3A-C and Table 1), similar to the phenotype observed upon silencing activity of the *Pdf*+ neurons via TNT. After eight days in DD, experimental flies showed a 3.3-hour phase delay in their activity peak (Fig. 3D-E). Under a light dark-dark cycle, *SLP316>NaChBac* flies showed a significant reduction in daytime activity (Suppl. Fig. 3A-D).

**Fig. 3:**
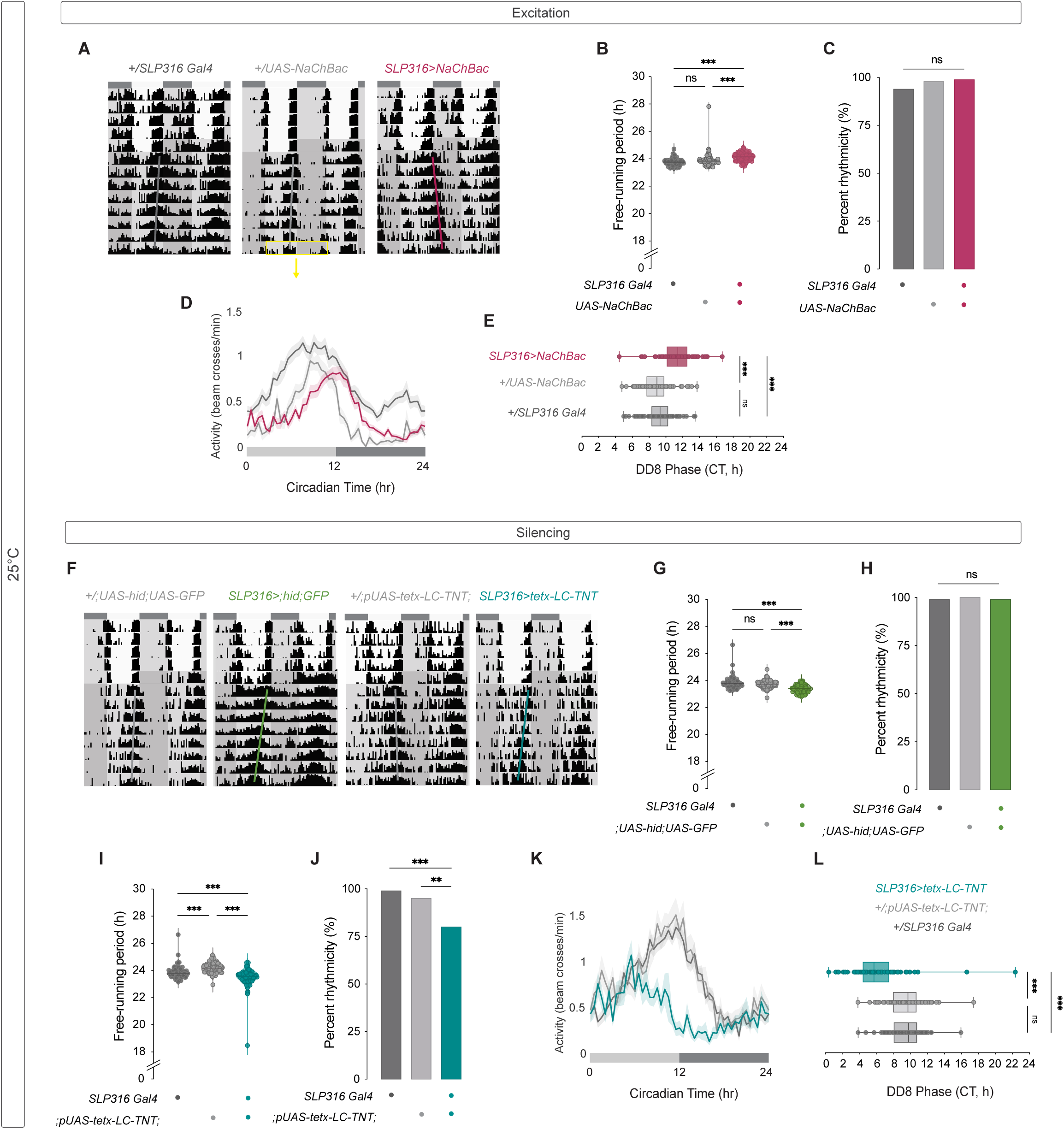
Manipulation of the SLP316s modulates free-running period length. (A) Representative double-plotted actograms, (B) free-running period calculated using the Chi-square periodogram, and (C) percentage of rhythmic flies for control (*SLP316 split Gal4* and *UAS-NaChBac*) and experimental (*SLP316>NaChBac*) flies. (D) Activity plot on the eighth day of free running, indicated in (A) with yellow box. (E) Phase of activity from (D), calculated using a sine-fit of the waveform of activity in CLOCKLAB. (F) Representative actograms, (G) free-running period and (H) percentage of rhythmic flies for control (w*;UAS-hid;UAS-GFP* and w*;pUAS-tetx-LC-TNT;*) and experimental (*SLP316 >hid+GFP* or *SLP316>tetx-LC-TNT*) flies. (I) Free-running period, and (J) percentage of rhythmic flies for control (*SLP316 split Gal4* and *;pUAS-tetx-LC-TNT;*) and experimental *(SLP316>tetx-LC-TNT*) flies. (K) Activity plot and (L) phase of activity on the eighth day of free-running for control (*SLP316 split Gal4* and *;pUAS-tetx-LC-TNT;*) and experimental *(SLP316 >tetx-LC-TNT*) flies. Actograms show 5 days of LD followed by 9 days of DD. Flies were raised and activity-rest behavior recorded at 25°C. Rhythmic power was calculated using the Chi-square periodogram in ClockLab using 30-minute activity bins, and flies with a rhythmic power of 10 or greater were classified as rhythmic. Free-running period was calculated for rhythmic flies using the Chi-square periodogram routine of ClockLab using 1-minute activity bins. Statistical comparisons were conducted using the Kruskal-Wallis test followed by Dunn’s multiple comparisons test for free-running period and phase of activity, and using Fischer’s exact test for the number of rhythmic and arrhythmic flies. Data are plotted from three replicate experiments for each genotype.

**Table 1.** Summary of free running activity rhythms. Activity analysis of the above genotypes at either 25°C or 18°C. Light conditions for each experiment were 12:12 LD-DD. For each genotype the number of flies (n) is listed above. All flies were raised at 25°C. ClockLab’s χ-square periodogram analysis was used to analyze rhythmicity, rhythmic power, and free-running period for each genotype. The percent rhythmicity, along with the number of rhythmic flies (nR), the period in hours with the SEM, and the rhythmic power with the SEM are indicated. Arrhythmic flies were not considered in the analysis. Statistical analysis is included in Table S1.

| Temperature: 25°C |  |  |  |  |  |
| --- | --- | --- | --- | --- | --- |
| Genotype | Sex | Number of Flies (n) | % Rhythmicity (nR) | Period (h) ± SEM | Rhythmic Power ± SEM |
| *;Pdf-Red,Pdf-Gal4;* | M | 90 | 100 (90) | 24.57 ± 0.04 | 148.3 ± 6.06 |
| *;pUAS-tetx-LC-TNT;* | M | 113 | 96.46 (109) | 24.11 ± 0.04 | 91.38 ± 4.75 |
| *;Pdf-Red,Pdf-Gal4; > ;tetx-LC-TNT;* | M | 76 | 78.95 (60) | 24.99 ± 0.11 | 64.58 ± 5.45 |
| *SLP316 split Gal4* | M | 170 | 95.88 (163) | 23.82 ± 0.03 | 117.4 ± 4.14 |
| *NaChBac* | M | 73 | 98.63 (72) | 23.91 ± 0.06 | 172.0 ± 5.82 |
| *SLP316 > NaChBac* | M | 69 | 98.55 (68) | 24.13 ± 0.04 | 171.0 ± 5.30 |
| *;UAS-hid;UAS-GFP* | M | 76 | 100 (76) | 23.73 ± 0.03 | 167.9 ± 5.39 |
| *SLP316 > ;hid;GFP* | M | 68 | 98.80 (82) | 23.38 ± 0.04 | 144.5 ± 6.98 |
| *SLP316 > ;tetx-LC-TNT;* | M | 85 | 80.0 (68) | 23.47 ± 0.09 | 41.02 ± 3.46 |
| *SLP316 split Gal4* | F | 31 | 80.65 (25) | 24.41 ± 0.12 | 89.78 ± 11.06 |
| *;UAS-hid;UAS-GFP* | F | 32 | 100 (32) | 24.39 ± 0.06 | 174.5 ± 8.34 |
| *SLP316 > ;hid;GFP* | F | 32 | 100 (32) | 24.26 ± 0.26 | 169.1 ± 11.20 |
| *;pUAS-tetx-LC-TNT;* | F | 29 | 93.10 (27) | 25.14 ± 0.34 | 117.4 ± 10.96 |
| *SLP316 > ;tetx-LC-TNT;* | F | 31 | 96.77 (30) | 24.25 ± 0.07 | 131.1 ± 7.69 |
| Temperature: 18°C |  |  |  |  |  |
| Genotype | Sex | Number of Flies (n) | % Rhythmicity (nR) | Period (h) ± SEM | Rhythmic Power ± SEM |
| *SLP316 split Gal4* | M | 160 | 97.50 (156) | 23.86 ± 0.11 | 81.45 ± 2.87 |
| *NaChBac* | M | 32 | 100 (32) | 23.72 ± 0.06 | 100.7 ± 5.74 |
| *SLP316 > NaChBac* | M | 32 | 100 (32) | 23.70 ± 0.07 | 106.5 ± 6.93 |
| *;UAS-hid;UAS-GFP* | M | 94 | 100 (94) | 23.78 ± 0.03 | 121.5 ± 4.91 |
| *SLP316 > ;hid;GFP* | M | 92 | 89.13 (82) | 23.53 ± 0.18 | 67.05 ± 4.41 |
| *;pUAS-tetx-LC-TNT;* | M | 94 | 79.79 (75) | 23.83 ± 0.18 | 49.49 ± 3.57 |
| *SLP316 > ;tetx-LC-TNT;* | M | 95 | 68.42 (65) | 23.45 ± 0.17 | 36.00 ± 2.84 |
| *Pdf01* | M | 31 | 93.55 (29) | 23.88 ± 0.04 | 63.02 ± 6.584 |
| *SLP316 split Gal4* | F | 63 | 84.13 (53) | 24.12 ± 0.18 | 79.16 ± 6.03 |
| *;UAS-hid;UAS-GFP* | F | 25 | 84.00(21) | 24.15 ± 0.13 | 66.93 ± 7.93 |
| *SLP316 > ;hid;GFP* | F | 26 | 92.31 (24) | 23.54 ± 0.56 | 84.16 ± 9.95 |
| *;pUAS-tetx-LC-TNT;* | F | 64 | 93.75 (60) | 24.29 ± 0.06 | 103.9 ± 6.31 |
| *SLP316 > ;tetx-LC-TNT;* | F | 58 | 91.38 (53) | 23.75 ± 0.09 | 82.40 ± 5.64 |

Next, we expressed *hid* under the SLP316 driver. Under DD conditions, *SLP316*>*hid* flies had a short free running period (Fig 3F-G). The overall number of rhythmic flies and their rhythmic power were not affected (Fig 3H and Table 1). A phase advance in activity can be detected on day 8 under DD conditions (Suppl. Fig. 3P-Q). *SLP316>hid* females also show a shortening in free-running period and no effect on the overall number of rhythmic flies (Suppl. Fig. 4A-B and E). Therefore, elimination of most SLP316 cells and *NaChBac*-mediated hyperexcitation of these cells led to opposite phenotypes in free-running period of activity rhythms.

To determine whether synaptic activity of the SLP316 cells mediates the effects on period we expressed TNT (34). This manipulation led to a shortened period (Fig 3F, I and Table 1). We also observed a small but significant reduction in the percentage of rhythmic flies and a phase advance at DD 8 of ∼3.2 hours (Fig. 3J-L). In *SLP316>TNT* females, the reduction in free-running period was not significantly different from that of controls (Suppl. Fig. 4C-E). Under LD, both *hid* and *TNT* expression in SLP neurons resulted in a reduction in daytime activity (Suppl. Fig. 3G, L). Shortening of free-running period is typically associated with an early phase of the evening peak of activity under a light-dark cycle (LD), such as in *Pdf*^01^ mutants or flies in which *Shaggy* or *Doubletime* (DBT^S^) are expressed in the sLN_v_s (57, 58). However, LD activity profiles of *SLP316>hid* and *SLP316>TNT* flies indicate that these flies could still anticipate the lights-on transition and did not have an early phase of the evening peak (Suppl. Fig. 3F,I,K and N). The evening peak area in experimental *SLP316>hid* and *SLP316>TNT* males was significantly lower than the controls, indicating lower activity at this time of day (Suppl. Fig.3J,O).

The only behavioral rhythm that is mediated by the LN_v_s but does not rely on PDF is adult emergence from the pupal case (eclosion) (59, 60). The larval LNs, which survive metamorphosis and become the adult sLN_v_s (61), communicate with the PTTH neurons to regulate eclosion rhythms (59). In pharate adults, the sLN_v_s inhibit PTTH neuron Ca^2+^ activity via short neuropeptide F (sNPF), not PDF, and the PTTH neurons appear to be post-synaptic targets of the sLN_v_s based on transsynaptic tracing experiments (59). In addition, PTTH neurons express sNPF and not PDF receptors (59). To determine if manipulations of SLP316 neurons affected eclosion rhythms, we conducted population experiments in which these cells were eliminated via expression of *hid*. We did not detect eclosion rhythm phenotypes in experimental *SLP316>hid* flies (Fig. 4A-B). Taken together, these results show that manipulations of the SLP316 neurons affect free-running rhythms in activity but not eclosion rhythms.

**Fig. 4:**
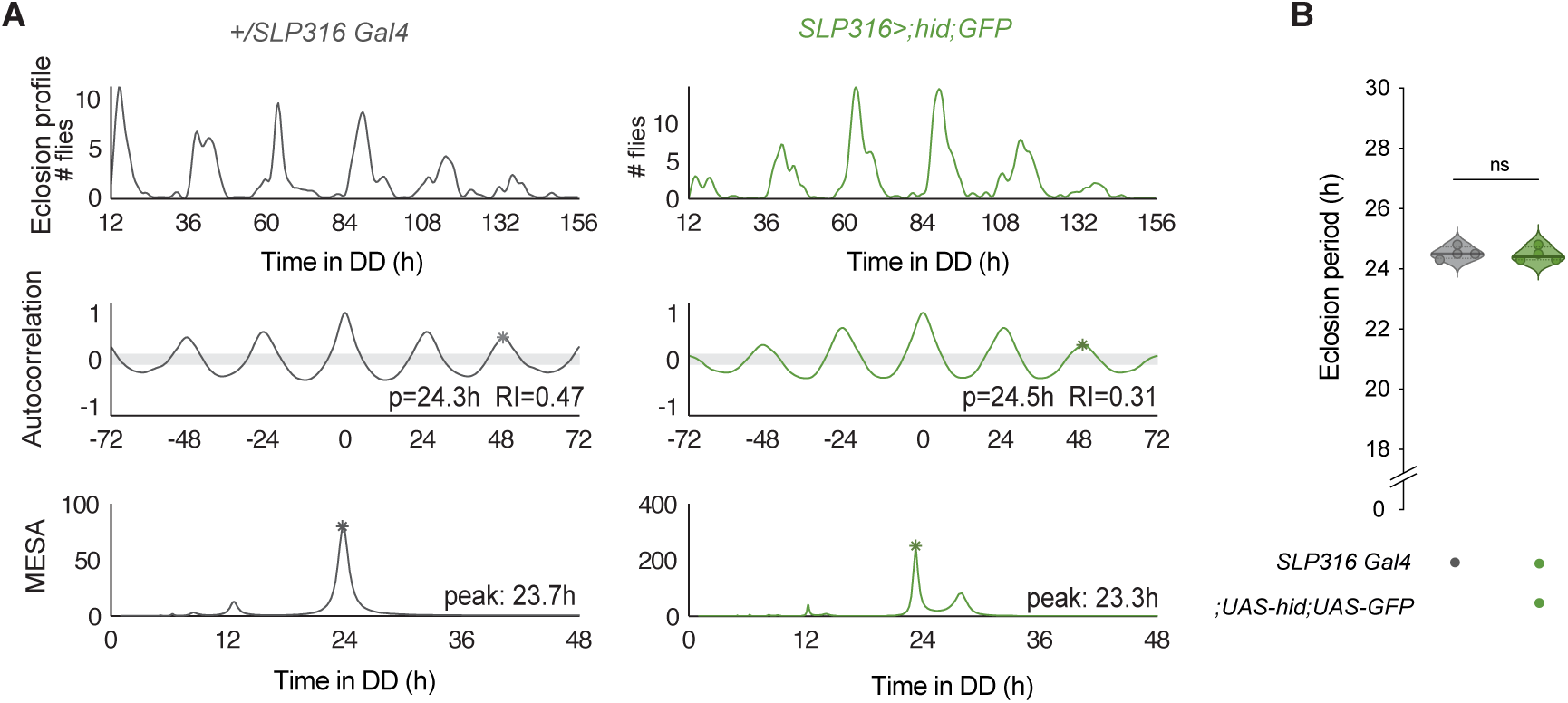
Silencing or eliminating SLP316 neurons does not affect the eclosion rhythm. (A) Records showing the time course of emergence of flies under DD conditions (left), their corresponding autocorrelation analysis (middle), and periodicities (right). (B) Period of eclosion of populations of flies of the indicated genotypes.

### 4. Organization of activity under cold days is disrupted upon SLP316 silencing

Since a subset of the DN1p class of clock neurons are major postsynaptic partners of the SLP316 cells, we investigated the effect of manipulating these cells under low temperature light-dark (LD) conditions, where the DN1ps appear to play a more prominent role in shaping activity (45). Only under conditions with low light intensity, low temperature, or low or absent PDF can rescuing the molecular clock in a subset of glutamatergic DN1ps promote an evening peak of activity (44, 45).

Low temperatures lead to a reorganization of activity in wild-type flies as they adapt to the shorter days of winter. Under low temperature conditions (18°C) and light-dark cycles, the distance between the morning and evening peaks is reduced, primarily due to a substantially earlier phase of the evening peak, which decreases the phase angle of entrainment (42). We exposed flies in which we eliminated or silenced SLP316 cells to LD cycles under cold temperatures. *SLP316>hid* flies show significantly reduced daytime activity and increased daytime sleep compared to controls (Fig. 5A-D). Similar phenotypes, both in terms of daytime activity and daytime sleep, were observed in experimental *SLP316>TNT* flies (Fig. 5E-H). Neither nocturnal activity nor sleep were affected (Suppl. Fig. 5A-D).

**Fig. 5:**
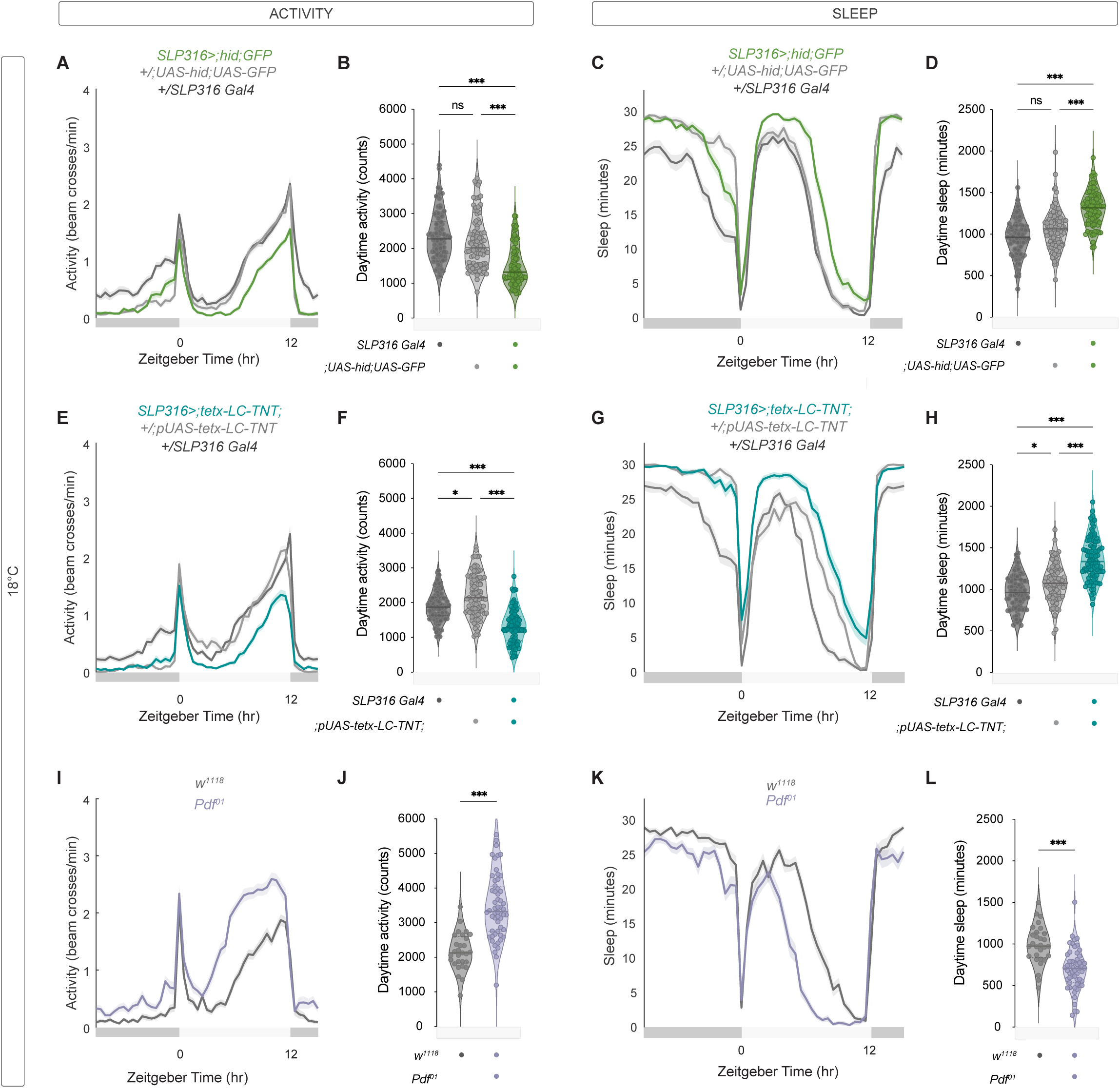
*Pdf* and SLP316 disruption have opposite effects on daytime activity and sleep quantities under cold temperature. (A-D) Activity and sleep phenotypes of *SLP316 >hid+GFP flies* during three days of 12:12 LD at 18°C. Representative activity (A) and sleep (C) plots averaged for (*SLP316 split Gal4* and *w;UAS-hid;UAS-GFP*) and experimental (*SLP316 >hid+GFP*) flies. Daytime (ZT0-12) (B) activity counts and total sleep amount (D). (E-H) Activity and sleep phenotypes of *SLP316 >;tetx-LC-TNT* flies and parental controls. Representative activity (E) and sleep (G) plots, and their respective quantifications (F, H). (I-l) Activity and sleep phenotypes of for wild-type *w^1118^* and *pdf^01^* flies. Representative activity (I) and sleep (K) plots, and total daytime activity counts (J) and sleep amount (L). Flies were raised at 25°C and activity-rest behavior was recorded at 18°C. Statistical comparisons were conducted using the Kruskal-Wallis test followed by Dunn’s multiple comparisons test for comparisons between three genotypes in (B), (D), (F) and (H), and using the Mann-Whitney test for comparisons between two genotypes in (J) and (L). Data plotted are from two replicate experiments for (B), (D), (F) and (H), and three replicate experiments for each genotype in (J) and (L).

Under free-running conditions, manipulations of SLP316 neuron activity and the loss of *Pdf* result in distinct phenotypes. The *Pdf*^01^ mutant is predominantly arrhythmic (18). However, the *Pdf*^01^flies that remained rhythmic exhibited a shortened free-running period, similar to the effect observed after silencing SLP316 neurons (18). Under light-dark (LD) conditions, manipulations of SLP316 neurons did not cause the expression of the classic phenotypes associated with the loss of *Pdf*, such as a lack of morning anticipation or an advanced evening peak (Suppl. Fig. 3). The advanced evening peak is thought to result from a phase-delay effect of PDF on evening cells: in the absence of PDF, neuronal activity in the evening cells show a large phase advance (24). PDF levels are temperature-sensitive, increasing during hot, summer-like days and decreasing under colder conditions (62).

We exposed *Pdf*^01^flies to the same LD 18°C cycles and observed a significant advance in the evening peak and a 1.5-fold increase in daytime activity (Fig. 5I-J). This indicates that the early peak observed in wild-type flies under LD (42) is not solely attributable to the reduction in PDF under lower temperatures, since flies that lack PDF show a pronounced shift in evening activity. *Pdf* mutants show higher levels of daytime sleep in 12:12 LD cycles at 25°C (63); however, the pronounced increase in evening activity under LD 18°C is accompanied by an overall decrease in daytime sleep (Fig. 5K-L). The onset of activity was earlier, and activity levels remained high until the lights-off transition. After the lights-off transition, activity and sleep levels between control and *Pdf*^01^ flies converge (Fig. 5I,K and Suppl. Fig. 6E-F). These findings indicate that under cold-day conditions, the loss of *Pdf* and the silencing or elimination of SLP316 neurons results in the expression of opposite behavioral phenotypes.

### 5. Silencing SLP316 neurons increases daytime sleep

Under standard light-dark conditions at 25°C, *Pdf* mutants show increased sleep during the day, as well as during the subjective day under constant conditions (63). However, the effect of *Pdf* loss on sleep appears to be dependent on temperature: indeed, these flies show increased sleep at 25°C and decreased sleep at 18°C (Fig. 5L). We analyzed the sleep profiles of flies expressing *hid* in the SLP316 cells under a LD cycle at 25°C and observed a significant increase in daytime sleep (Fig. 6A-B). (These results were obtained using the standard quantification based on periods of 5 minutes of inactivity (64, 65)). A similar phenotype was observed in *SLP316*>*TNT* flies (Fig. 6E-F). Neither of those manipulations affected nighttime sleep (Suppl. Fig 6). These findings allow us to conclude that the effects on sleep are not unique to low temperature cycles but, in contrast to PDF’s temperature dependent roles, reflect a feature of the SLP316 neurons at both low and high temperature.

**Fig. 6:**
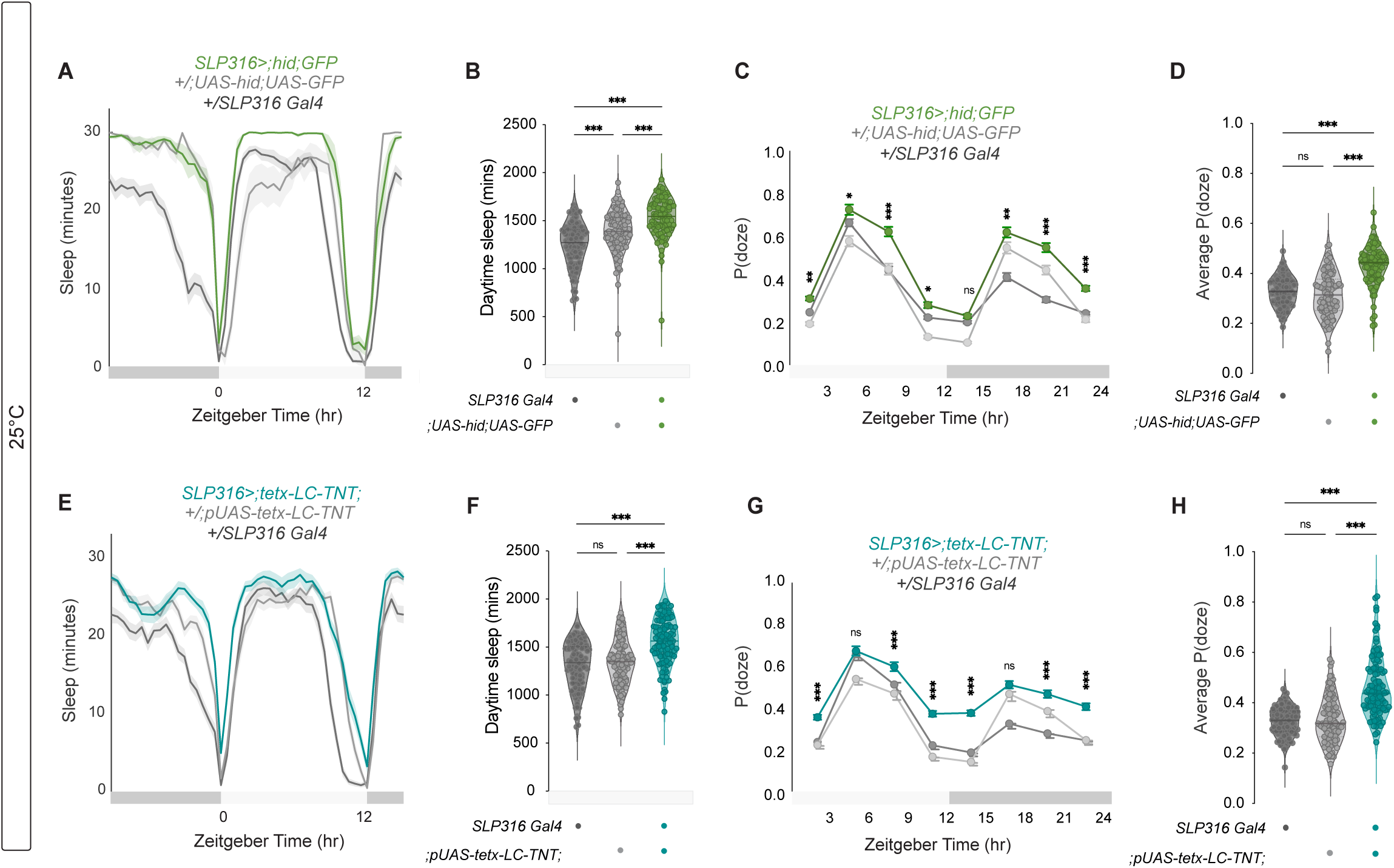
Silencing or eliminating SLP316 neurons increases the amount of daytime sleep. (A) Representative sleep plot averaged from three days of 12:12 LD at 25°C for control (*SLP316 split Gal4* and *;UAS-hid;UAS-GFP*) and experimental (*SLP316 >;hid;GFP*) flies. (B) Total daytime sleep over three days for each genotype. (C) Representative sleep plot and (D) total daytime sleep for control (*SLP316-Gal4* and *;pUAS-tetx-LC-TNT;*) and experimental (*SLP316 >tetx-LC-TNT*) flies. (E) P(doze) from the same three days of LD, plotted in three-hour for either control shown for controls, *SLP316 split Gal4* and *;UAS-hid;UAS-GFP*), and experimental (*SLP316 >;hid;GFP*) flies. (F) Average P(doze) throughout entire 3-day period. (G) P(doze) plot and (H) average P(doze) for control (*SLP316-Gal4* and *;pUAS-tetx-LC-TNT;*) and experimental (*SLP316 >tetx-LC-TNT*) flies under same conditions. Sleep was quantified in PHASE based on periods of inactivity of 5 minutes or more and plotted in (A) and (C) as minutes of sleep per 30-minute bin. P(doze) was quantified in R using routines from the Griffith Laboratory (66). Statistical comparisons were conducted using the Kruskal-Wallis test followed by Dunn’s multiple comparisons test. Data is plotted from four replicate experiments.

To determine the underlying cause for this increase in sleep, we analyzed P(doze) and P(wake), the probabilities that an inactive fly will start moving or an active fly will stop moving, respectively, between 1-minute activity bins. P(doze) corresponds to sleep pressure, and P(wake) corresponds to sleep depth; thus, these conditional probabilities can offer insight into the biological processes driving changes in sleep (66). P(doze) was higher throughout most of the day and night for both *SLP316>hid* and *SLP316>TNT* animals, especially during the evening (ZT6-12) and morning (ZT18-3), when flies are most active (Fig. 6 C,D and G,H). The average P(wake) was not significantly affected in either manipulation, although *SLP316>hid* flies showed a minor reduction in P(wake) from ZT9-12 (Suppl. Fig. 7A-D). This increase in P(doze) suggests that the increased sleep observed when the SLP316s are silenced was due to increased sleep pressure, especially at times when flies should be most active. We conclude that silencing SLP316 neurons leads to an increase in daytime sleep under both 25°C and low temperature conditions, and that this phenotype is not dependent on PDF since *Pdf* manipulations lead to opposite effects.

## Discussion

The recent availability of connectomics data allows the identification of neuronal connections that have not been described using other approaches. We used openly available *Drosophila* connectomes to identify the SLP316 neurons, which are the main post-synaptic partners of the sLN_v_s, the key pacemaker neurons in the *Drosophila* clock network. We found that genetic manipulations of the SLP316 neurons led to changes in free-running rhythms: silencing their activity via TNT or eliminating them via *hid* leads to a period shortening, whereas hyperexciting them via *NaChBac* led to a period lengthening. Another important behavioral output of the sLN_v_s, eclosion rhythm, was not affected by genetic elimination of SLP316 neurons. Under light dark cycles, silencing SLP316s resulted in a reduction in daytime activity and an increase in sleep, under both 25°C and low temperature conditions. Neither the free-running nor the light-dark (LD) phenotypes observed following SLP316 manipulations resembled those caused by the loss of PDF, a well-known output of the sLN_v_s, which facilitates communication with the rest of the circadian clock network.

Our study identified a non-clock group of neurons that affects circadian rhythms and contributes to our understanding of neuronal connectivity within the clock network. To date, the best characterized output of the sLN_v_s is PDF (18, 67), a neuropeptide that accumulates--and is also released (33)-rhythmically from their dorsal termini and soma, likely independently of classical transmitter release from these cells (32). The sLN_v_s also express sNPF (28), and there is evidence that they also use glycine to communicate with postsynaptic targets (31). Based on the availability of neurotransmitter use information, the flywire EM prediction for the sLN_v_s is acetylcholine (50), but the likelihood is relatively low. If the sLN_v_s contact the SLPs via glycine, they would likely inhibit them, which is consistent with the observation that inhibiting synaptic release from *Pdf* neurons via TNT and exciting the SLP316 neurons via *NaChBac* both lead to period lengthening. Functional connectivity experiments between the sLN_v_s and the SLPs would be needed to determine the nature of this connection.

Given their prominent role in the clock network, several studies have examined neuronal targets of the sLN_v_s that would contribute to the propagation of rhythmic signals throughout the brain. PDF receptors (*PdfR*) are expressed by a wide range of neurons distributed throughout the brain, including much of the clock network (6), and PDFR-expressing cells respond to sLN_v_ stimulation (68). PTTH neurons, which are downstream of sLN_v_s, express sNPF receptors, are inhibited by the sLN_v_s via sNPF to regulate eclosion rhythms (59). Connections between *Pdf*+ cells and non-clock cells that regulate rhythm amplitude were detected via GRASP (69). More recently, the Blau laboratory used the transsynaptic tracer trans-tango to identify neuronal populations downstream of the sLN_v_s and reported that they form connections with a subset of DN1p clock neurons that express the neuropeptide CNMa and with mushroom body Kenyon cells (70). Strong monosynaptic connections between the sLN_v_s and the DN1ps cannot be detected in the connectomes (16, 49), but it is possible that those synapses were missed because the sLN_v_s show daily structural plasticity rhythms (71), as do the VIP expressing neurons in the SCN (72). Truncation of the dorsal termini of the sLN_v_s, which are the sites of synaptic output and undergo clock-controlled plasticity, does not affect free-running rhythms (38, 73), suggesting that PDF release is independent of the arborization rhythms. In addition, silencing synaptic transmission in *Pdf*+ neurons via TNT does not lead to the loss of behavioral rhythms (36), nor does downregulation of *bruchpilot* (*brp*) or other active zone proteins (32). Taken together, these results suggest that unlike PDF release, the synaptic connections that the sLN_v_s form with downstream neurons are not required for maintaining activity-rest rhythms under free running conditions. However, those connections play a role in regulating the free-running period of those rhythms.

The sLN_v_s appear to contact the DN1p clock neurons both via PDF release (58) and synaptically through the SLP316s (Fig. 1). A functional clock in CRY-positive DN1ps, along with the presence of PDF, is sufficient for morning anticipation (44, 45, 74). Neither inhibition of synaptic transmission from sLN_v_ nor manipulation of the SLP316 cells appeared to affect morning anticipation, suggesting that this feature depends on only PDF signaling from the sLN_v_s. Rescuing the molecular clock in a glutamatergic subset of DN1ps can promote an evening peak of activity when PDF is reduced or absent (44, 62). The pronounced and very early peak of activity we observed in *Pdf^01^*flies under cold temperatures may at least in part be due to increased prominence of the activity of evening DN1ps in the absence of PDF (75). Another clock subset, DN1a, which is inhibited by cold temperature, is connected reciprocally to the sLN_v_s through the neuropeptides CCHamide1 and PDF (76, 77). Thus, a cold-temperature induced change in sLN_v_ activity could alter evening DN1p activity via the SLP316s, while simultaneous loss of PDF eliminates suppression of the DN1p-induced effects. These glutamatergic DN1ps are also sleep-promoting; activating them increases daytime sleep and arousal threshold at ZT6 and 10, whereas inhibiting them decreases both daytime and nighttime sleep (78). If the SLP316 neurons are glutamatergic as predicted in Flywire (49) and inhibitory, silencing them would be expected to increase the neuronal activity of the DN1ps and thus daytime sleep levels, in agreement with our results.

Studying how neurons use different signals to communicate with downstream targets is essential for understanding how networks of neurons produce behavior. The *Drosophila* sLN_v_s are an excellent model for studying neuronal basis of behavior given their well characterized role as key circadian pacemakers. This role relies on their release of the neuropeptide PDF as a critical output signal for synchrony within the network. Our study is the first to examine the role on rhythmic behavior of direct synaptic targets of the sLN_v_s based on connectomics data, a connection that is likely independent of PDF. Our results reveal that manipulations of the SLP316 cells, the primary postsynaptic partners of the sLNvs in the connectome, can affect a core circadian parameter, free-running period. This finding indicates that a non-clock neuronal cell type can modulate circadian timekeeping. We show that manipulations of the SLP316 cells and loss of PDF lead to different behavioral phenotypes, both under free running conditions and under standard and low temperature light-dark cycles. These two pathways of clock neuron output may add plasticity to the system and allow the circadian network to adjust to changing environmental conditions.

## Materials and Methods

### Fly lines and husbandry

Flies were reared on cornmeal-sucrose-yeast media under a 12hr:12hr Light:Dark (LD) cycle at 25 °C. The fly lines used in this study were: *SLP316 split-Gal4* (SS76489: w; 38E08-p65ADZp in attP40; 36B07-ZpGdbd in attP2), *w*; *Pdf-Red, Pdf-Gal4; w;;UAS-NaChBac, yw;UAS-hid/cyo;UAS-mCD8-GFP, w;UAS-TeTxLC;, w^1118^, w;;Pdf^01^*, and *w;* LexAop2-Syb.GFPN146I.TEVT173V LexAop2-QF2.V5.hSNAP25.HIVNES.Syx1A/CyO,20XUAS-B3R.PEST-UAS(B3RT.B2)BoNTA QUAS-mtdTomato-3xHA/TM6B, Tb1 (see Fly Lines and Reagents table for details).

We identified the SLP316 split-GAL4 line (SS76489) by screening candidate hemidriver combinations selected by comparing GAL4 driver expression patterns and the hemibrain reconstructions of SLP316. This work was done in collaboration with the Janelia Fly Light Project Team using established methods and the driver line is included in a recently published split-GAL4 collection (79).

### Immunohistochemistry

Flies were entrained to 12:12 LD cycles at 25°C and dissected at the specified times in cold Schneider’s *Drosophila* Medium. For all experiments, dissections were completed within 30 minutes. For anti-PER staining (Fig. 2D), brains were dissected at ZT23 on the fifth day under LD. Brains were fixed in 2% paraformaldehyde prepared in Schneider’s *Drosophila* Medium for 30 minutes at room temperature. Blocking was performed using 5% normal goat serum in PBS containing 0.3% Triton (PBS-TX) for 1 hour at room temperature. Samples were incubated with primary antibodies for 24 to 48 hours at 4°C, followed by several washes with PBS-TX under agitation. Secondary antibody cocktails were applied for 24 to 48 hours at 4°C, after which samples were washed again with PBS-TX. Antibodies (see table) were diluted in PBS-TX supplemented with 5% normal goat serum. Brains were mounted between coverslips and imaged at the Advanced Science Research Center (GC-CUNY) using an Olympus Fluoview 3000 laser-scanning confocal microscope (Olympus, Center Valley, PA). Brains for the BacTrace experiments were imaged using an Olympus/Evident Fluoview 4000 laser-scanning confocal microscope (Fernández lab, Indiana University Bloomington).

The LM images in Figure S2 are based on confocal stacks previously reported (79) and the original images are available online (https://splitgal4.janelia.org/cgi-bin/view_splitgal4_imagery.cgi?line=SS76489). The EM-LM overlay was generated using VVDviewer (https://zenodo.org/records/14468001). The other images show maximum intensity projections produced using Fiji (https://imagej.net/software/fiji/).

### Connectome analysis

The *Drosophila* Hemibrain (release v2.1) was accessed via neuprint (https://neuprint.janelia.org/). FlyWire data are based on FAFB version 783 (https://flywire.ai/)(49, 80). Hemibrain reconstructions are identified by their instance names in neuprint (e.g. ‘SLP316_R’) with individual cells distinguished here by an additional number (‘SLP316_R_1’).

### Behavior recording and analysis

Locomotor activity rhythms of adult male and virgin female flies were recorded using DAM2 *Drosophila* Activity Monitors. Three- to six-day old flies were placed individually in TriKinetics capillary tubes containing 2% agar-4% sucrose food at one end sealed with a cap, plugged with a small length of yarn, and loaded into the DAM2 monitors for locomotor activity recording. Flies were entrained to 12:12 LD cycles for at least five days and then released into constant darkness (DD) for at least nine days at a constant temperature of 18 or 25°C (as indicated in the figure legend for each experiment).

Free-running activity rhythms were analyzed with ClockLab software. ClockLab’s Chi-square periodogram function was used to analyze rhythmicity and rhythmic power in individual flies using 30-minute bins for activity and a confidence level of 0.01. The “Rhythmic Power” for each designated rhythmic fly was determined by subtracting the “Significance” value from the “Power” value associated with the predominant peak. Flies that did not exhibit a Rhythmic Power of 10 were categorized as “arrhythmic,” and their period and rhythmic power were not included in the analysis (81). ClockLab periodogram analysis was repeated for rhythmic flies using 1-minute bins to quantify free-running period. For each of the genotypes tested, only periodicities falling within the 14 to 34-hour range were taken into consideration. Phases on the eighth day of free-running were calculated using 1-minute bins and applying a Least Squares sine-wave fit in the ClockLab activity profile, excluding any phases with P>0.05. All relevant information about statistical analyses is included in Table S1.

Activity counts were collected in 1-minute bins, and averaged across the population and profiles of specific genotypes under LD were generated using PHASE (82). First, individual fly activity levels were normalized by establishing the average activity across all 30-minute bins over days 2-4 under LD. Population averages of this normalized activity were computed for each 30-minute bin and averaged into a single representative 24-hour day. Total daytime (ZT0-12) and nighttime (ZT12-24) activity counts from days 2-4 of LD were recorded. Representative sleep plots and amounts were similarly generated for LD days 2-4 by classifying 5-minute periods of inactivity as sleep, unless otherwise specified. Total daytime and nighttime sleep over the three-day period were also recorded. Morning and evening phase were quantified in PHASE using a filter frame length of 481 minutes and filter order of 2. P(wake) and P(doze) values (66). were averaged for the entire duration of LD days 2-4, or in three-hour windows and averaged across the same days.

### Adult emergence monitoring

Flies were raised at 25°C for 5 days, then transferred to 20°C and entrained under a LD 12:12 light:dark cycle. Resulting pupae were collected and fixed on eclosion plates with Elmer’s glue and mounted on Trikinetics eclosion monitors (Trikinetics, MA, USA). Emergence was then monitored for 7 days at 20°C under DD conditions and analyzed as previously described (83). Rhythmicity of eclosion profiles was analyzed using MATLAB (MathWorks, Inc., Natick) and the appropriate MATLAB toolbox (84). Using autocorrelation analyses, records were considered rhythmic if the rhythmicity index (RI) was 0.3, weakly rhythmic if 0.1<0.3, and arrhythmic if RI<0.1 (20).

### Fly lines and reagents

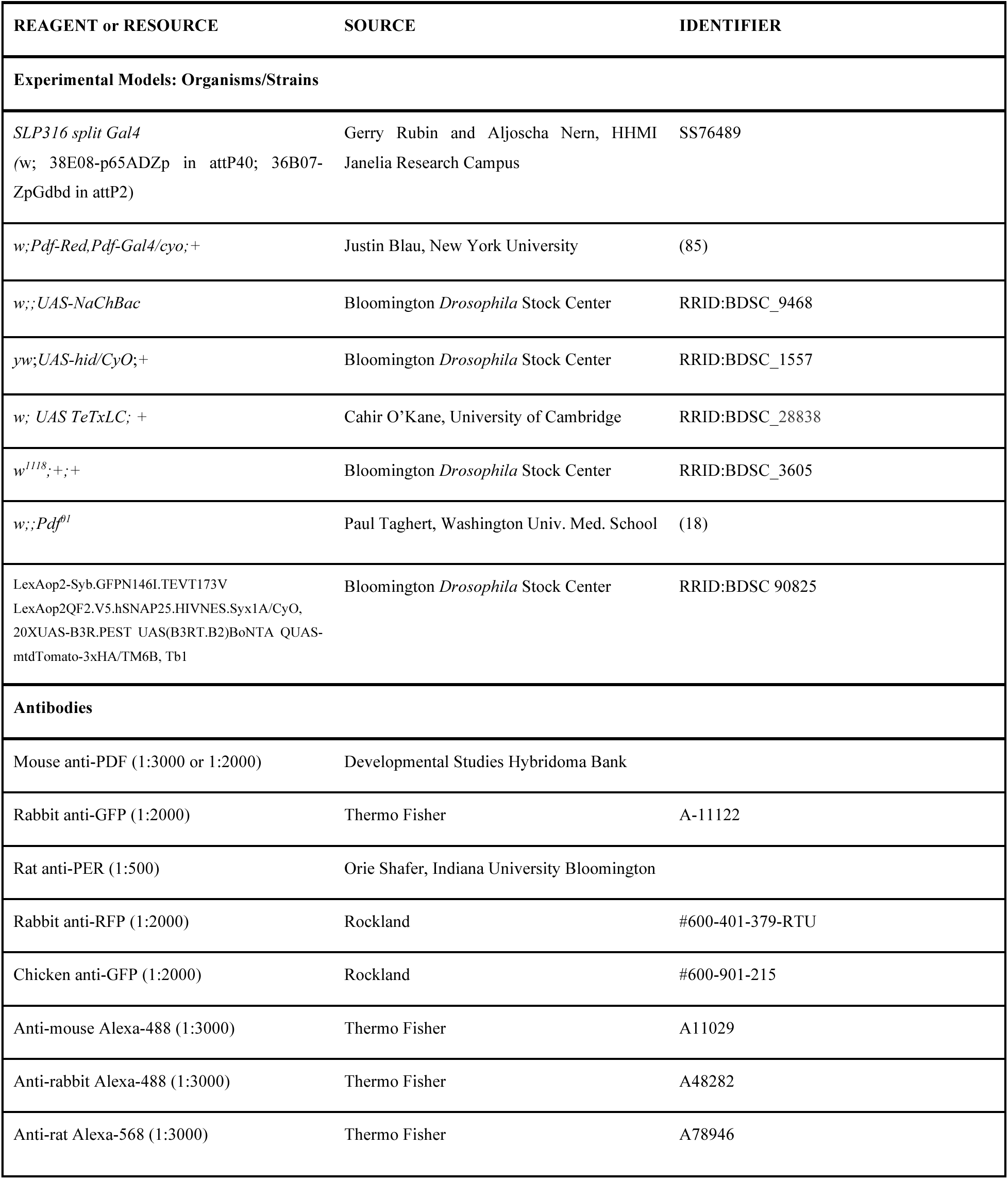
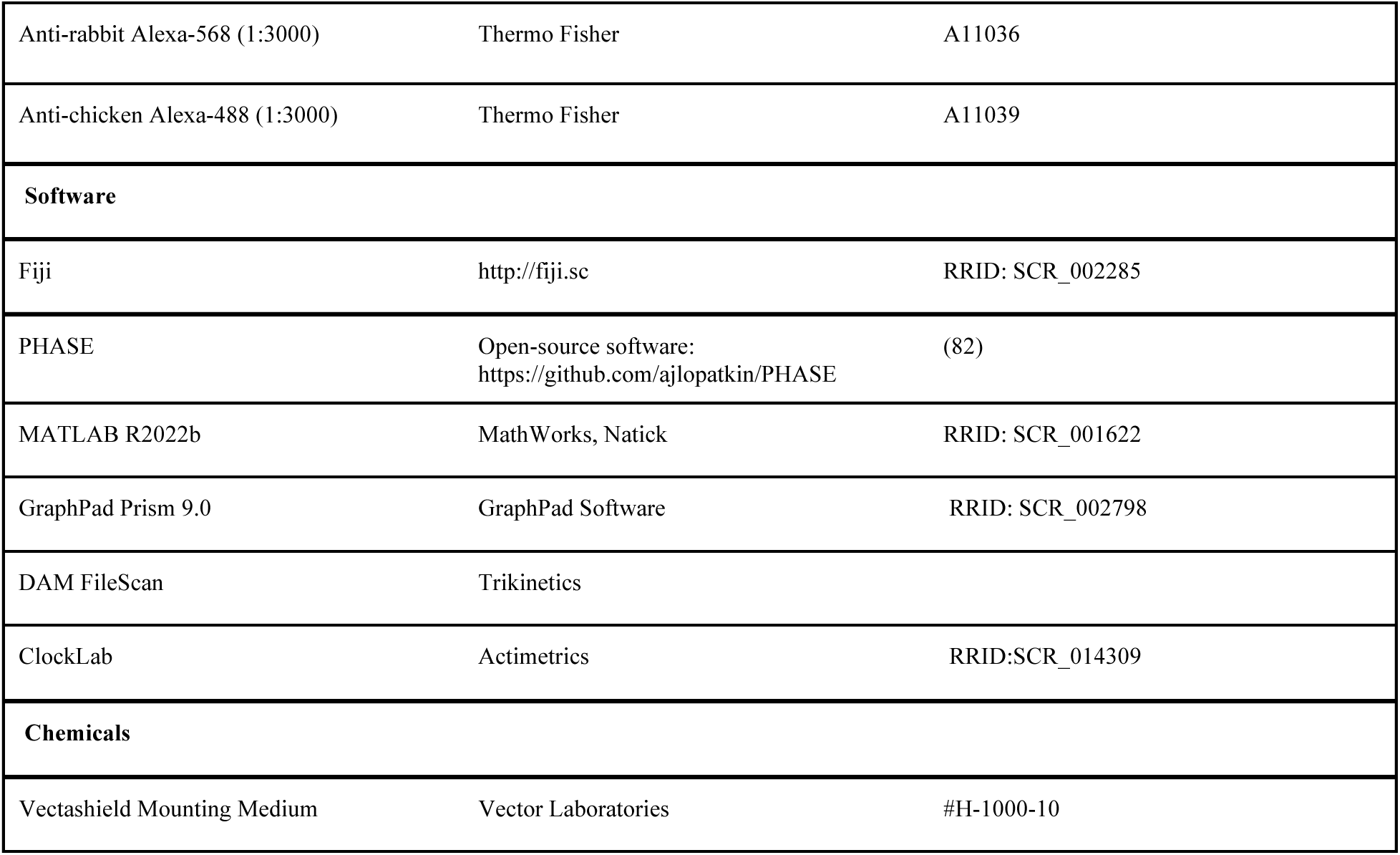

## Acknowledgments

We are grateful to Patrick Emery and members of the Fernández lab for helpful discussions and to Orie Shafer for comments on the manuscript, assistance on imaging, and providing access to the FV3000 microscope. We also thank the Janelia *FlyLight* Project Team for help with split-GAL4 screening and imaging, Justin Blau, Paul Taghert, and Cahir O’Kane for sharing fly lines, and Abhilash Lakshman for the code to obtain P(wake) and P(doze) values. Stocks obtained from the Bloomington *Drosophila* Stock Center (NIH P40OD018537) were used in this study. The mouse anti-PDF antibody was obtained from the Developmental Studies Hybridoma Bank, created by the NICHD of the NIH and maintained at The University of Iowa, Department of Biology, Iowa City, IA 52242, and the rat anti-PER antibody was donated by O. Shafer. This work was supported by a Beckman Foundation Scholarship to E.S-C, an NSF CAREER award (IOS-2239994) to M.P.F. and an NIH (R01NS118012) grant.

