## Supplementary Figures and Table S1 for "Synaptic Targets of Circadian Clock Neurons Influence Core Clock Parameters"

Fig. S1

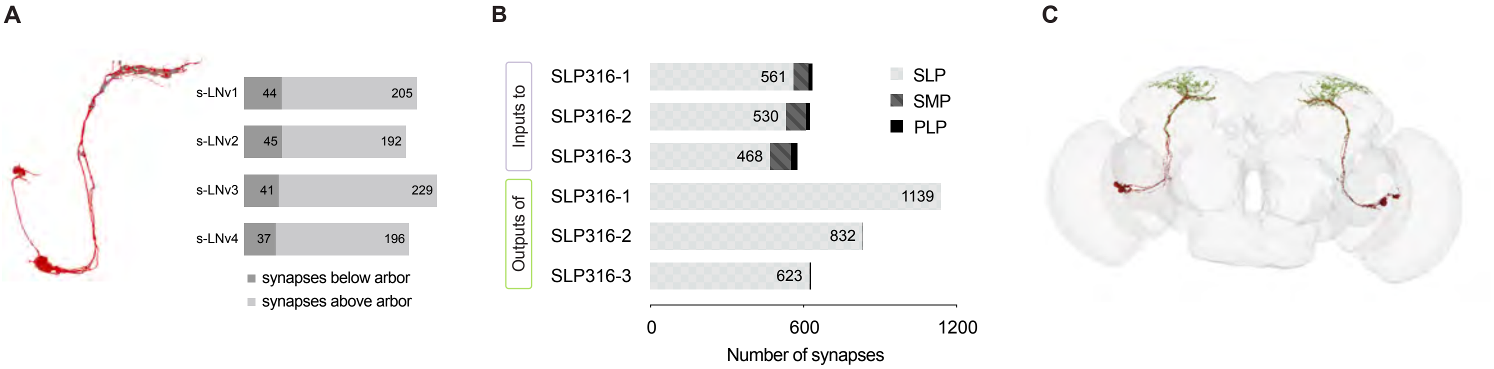
A
s-LNv1
44
205
s-LNv2
45
192
s-LNv3
41
229
s-LNv4
37
196
synapses below arbor
synapses above arbor
B
Inputs to
Outputs of
SLP316-1
561
SLP316-2
530
SLP316-3
468
SLP316-1
1139
SLP316-2
832
SLP316-3
623
SLP
SMP
PLP
0
600
1200
Number of synapses
C

Fig. S2

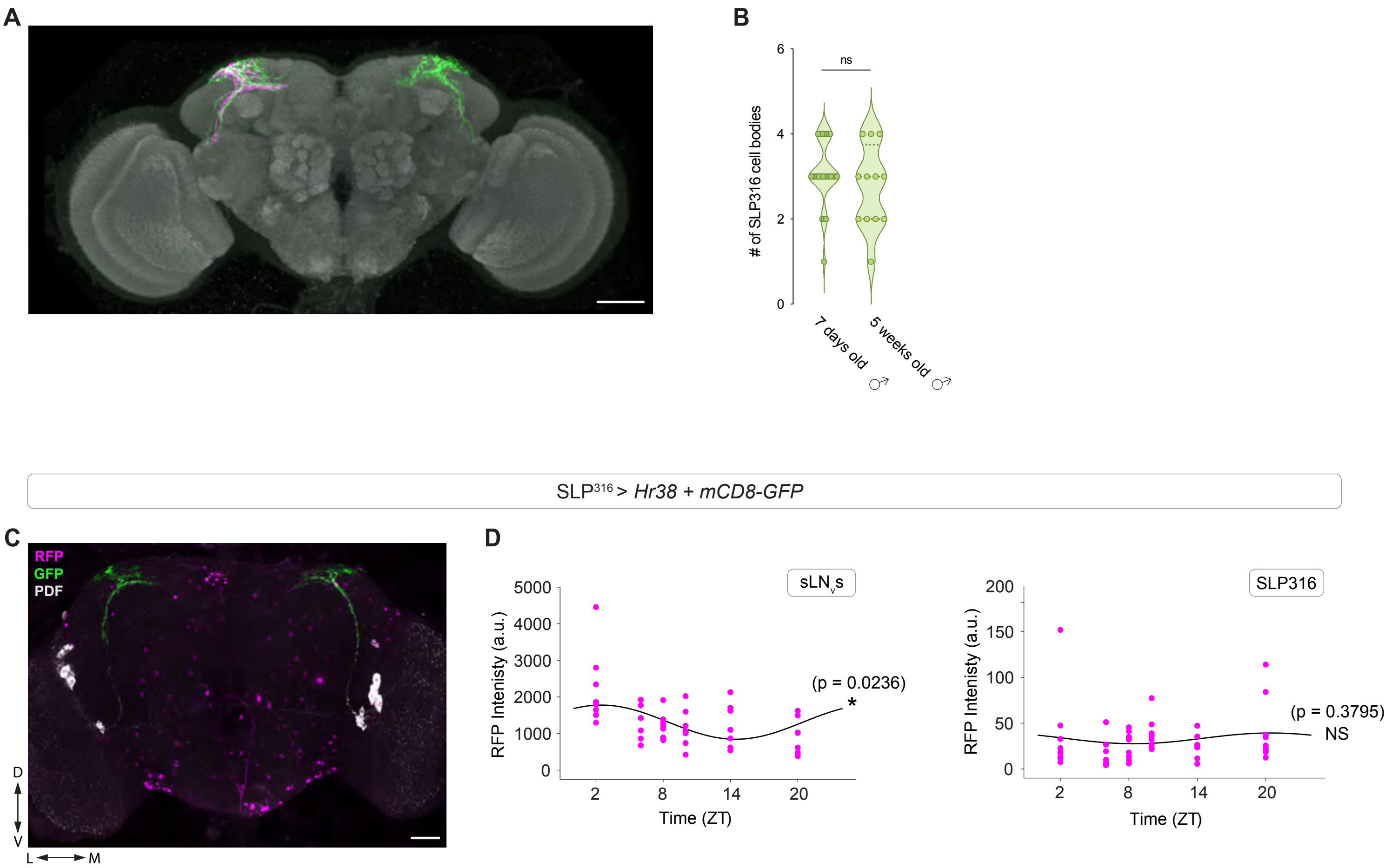

A
B
6
4
2
0
ns
### of SLP316 cell bodies
7 days old ♂
5 weeks old ♂
SLP316 > Hr38 + mCD8-GFP
C
RFP
GFP
PDF
D
V
L
M
D
sLNvs
RFP Intenisty (a.u.)
5000
4000
3000
2000
1000
0
2
8
14
20
Time (ZT)
(p = 0.0236)
*
SLP316
200
150
100
50
0
(p = 0.3795)
NS

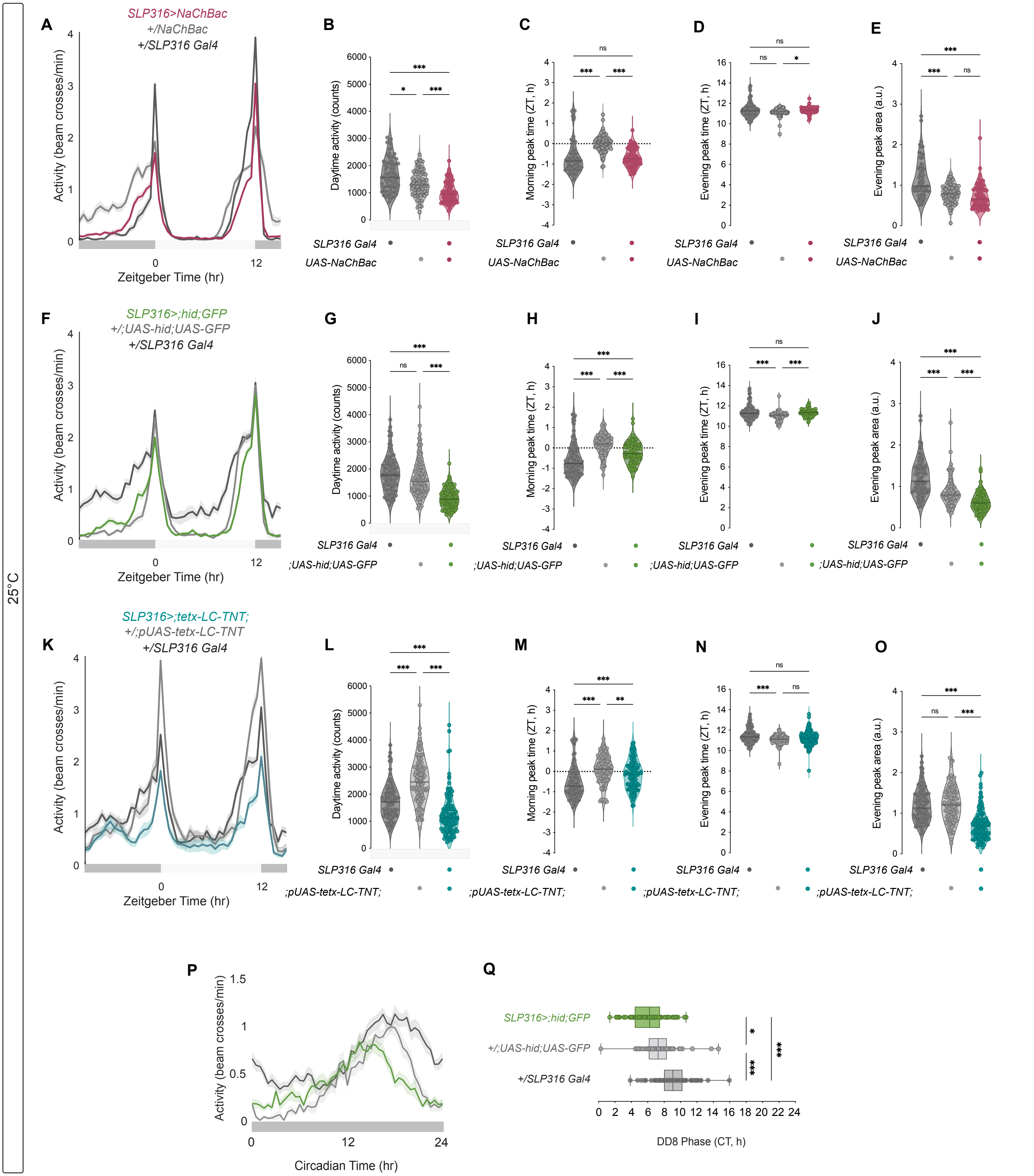
25°C
A
SLP316>NaChBac
+/NaChBac
+/SLP316 Gal4
Activity (beam crosses/min)
Zeitgeber Time (hr)
B
Daytime activity (counts)
C
Morning peak time (ZT, h)
D
Evening peak time (ZT, h)
E
Evening peak area (a.u.)
SLP316 Gal4
UAS-NaChBac
F
SLP316>;hid;GFP
+/;UAS-hid;UAS-GFP
+/SLP316 Gal4
G
H
I
J
;UAS-hid;UAS-GFP
K
SLP316>;tetx-LC-TNT;
+/;pUAS-tetx-LC-TNT
+/SLP316 Gal4
L
M
N
O
;pUAS-tetx-LC-TNT;
P
Circadian Time (hr)
Q
DD8 Phase (CT, h)
ns
*
**
***

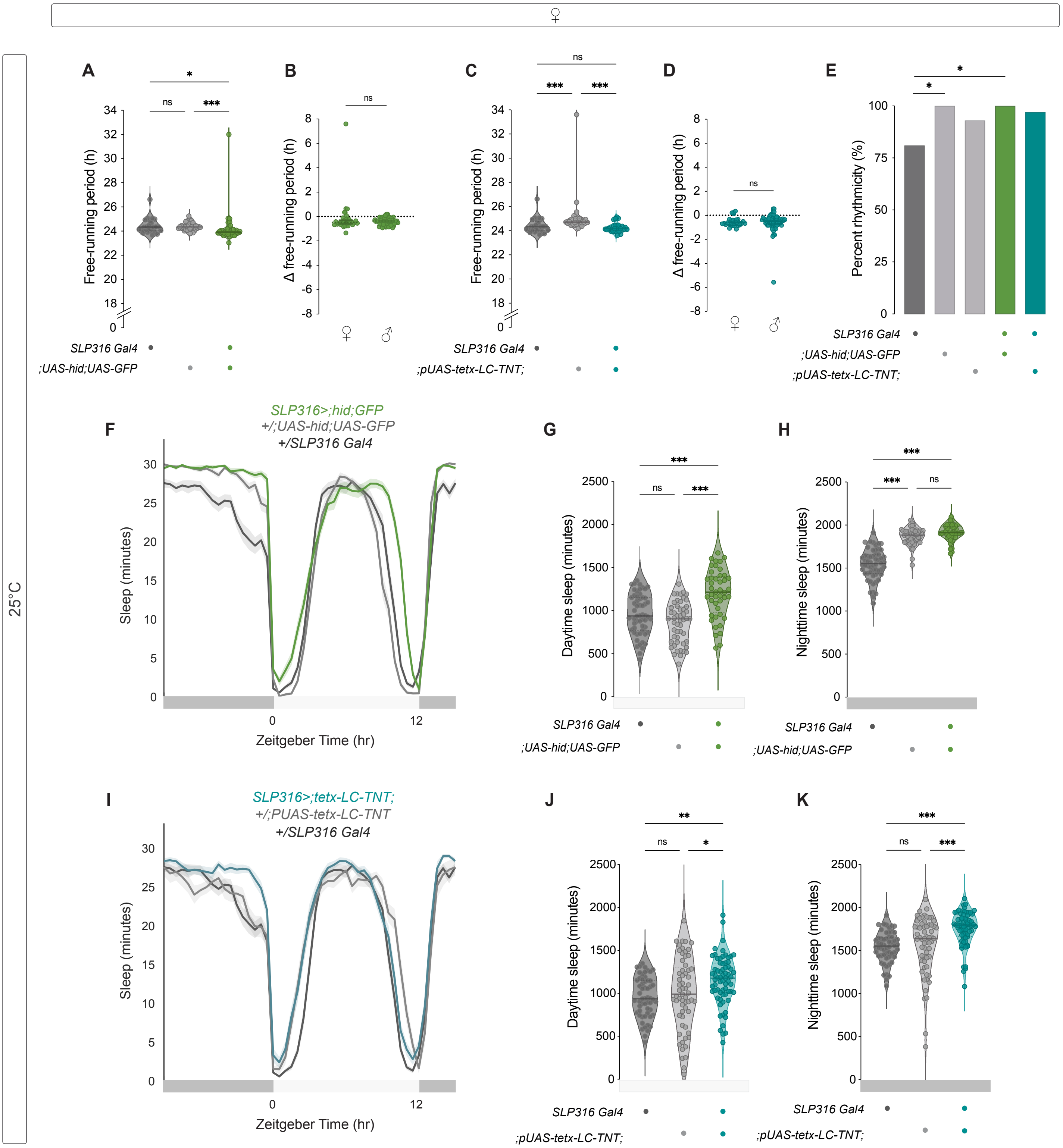
♀
25°C
A
Free-running period (h)
SLP316 Gal4
;UAS-hid;UAS-GFP
B
Δ free-running period (h)
C
;pUAS-tetx-LC-TNT;
D
E
Percent rhythmicity (%)
F
SLP316>;hid;GFP
+/;UAS-hid;UAS-GFP
+/SLP316 Gal4
Sleep (minutes)
Zeitgeber Time (hr)
G
Daytime sleep (minutes)
H
Nighttime sleep (minutes)
I
SLP316>;tetx-LC-TNT;
+/;PUAS-tetx-LC-TNT
J
K

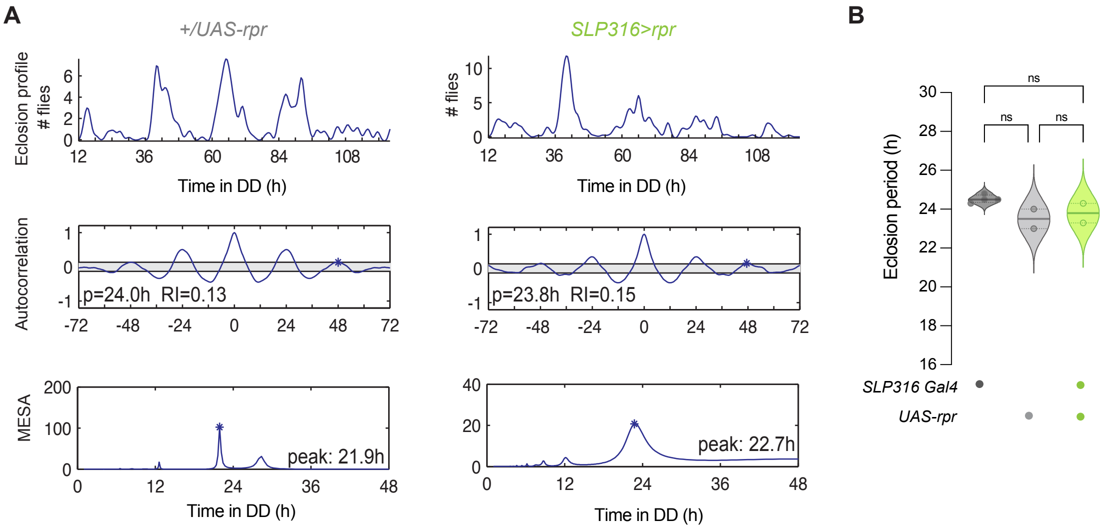
A
+/UAS-rpr
SLP316>rpr
Eclosion profile
### flies
Time in DD (h)
Autocorrelation
p=24.0h RI=0.13
p=23.8h RI=0.15
MESA
peak: 21.9h
peak: 22.7h
B
ns
Eclosion period (h)
SLP316 Gal4
UAS-rpr

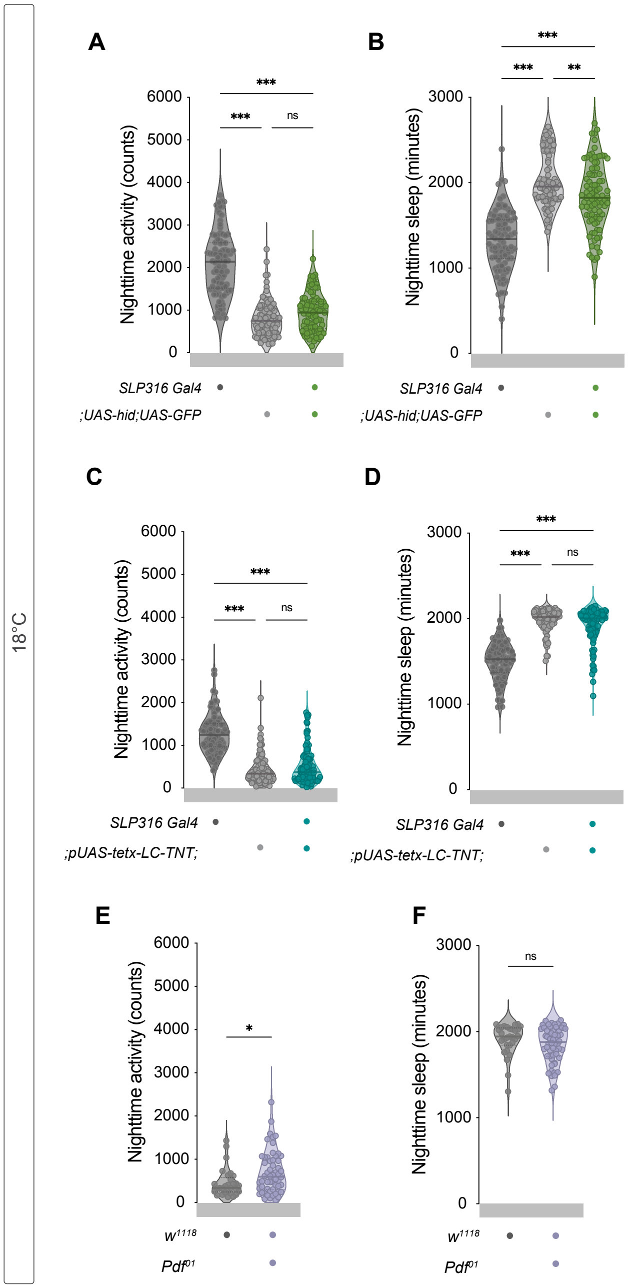
18°C
A
B
C
D
E
F
Nighttime activity (counts)
Nighttime sleep (minutes)
***
**
ns
*
SLP316 Gal4
;UAS-hid;UAS-GFP
;pUAS-tetx-LC-TNT;
w1118
Pdf01

Fig. S7

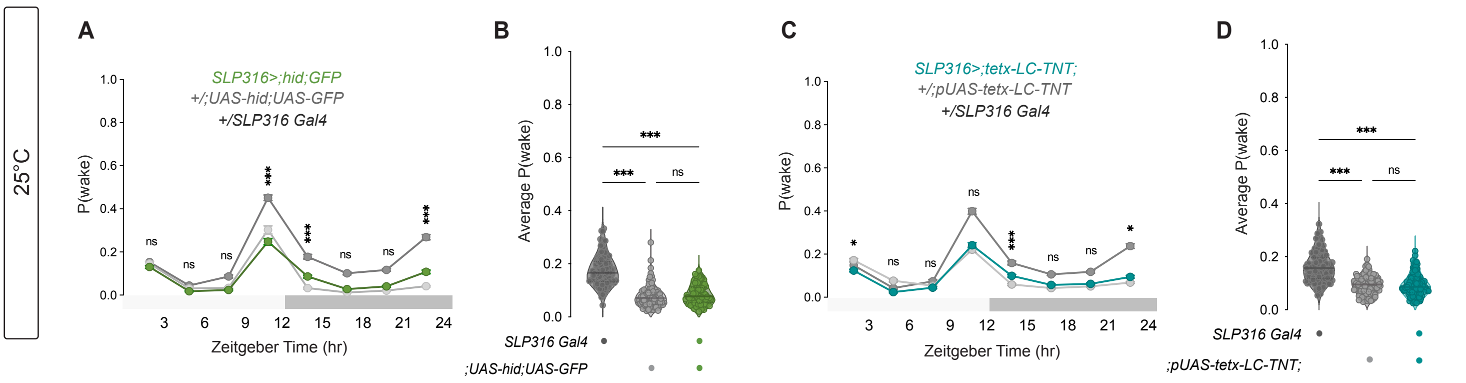
25°C
A
SLP316>;hid;GFP
+/;UAS-hid;UAS-GFP
+/SLP316 Gal4
P(wake)
Zeitgeber Time (hr)
B
Average P(wake)
SLP316 Gal4
;UAS-hid;UAS-GFP
C
SLP316>;tetx-LC-TNT;
+/;pUAS-tetx-LC-TNT
+/SLP316 Gal4
P(wake)
Zeitgeber Time (hr)
D
Average P(wake)
SLP316 Gal4
;pUAS-tetx-LC-TNT;

**Table S1.**
**Statistical**

| Figure | Quantification | Sex | Age/Sex 1 | Age/Sex 2 | Age/Sex 3 | Age/Sex 4 | Comparison | Test Used | P Value |
|---|---|---|---|---|---|---|---|---|---|
| 2 | 2C. SLP316 cell body count | M and F | 7 day old male | 7 day old female | 1 day old female | 1 day old male | 7 day old male vs 7 day old female | Tukey's Multiple Comparisons Test | 0.9996 (adjusted) |
| | | | | | | | 7 day old male vs 1 day old male | Tukey's Multiple Comparisons Test | 0.0048 (adjusted) |
| | | | | | | | 7 day old female vs 1 day old female | Tukey's Multiple Comparisons Test | 0.0087 (adjusted |
| | | | | | | | 1 day old male vs 1 day old male | Tukey's Multiple Comparisons Test | 0.9983 (adjusted) |

| Figure | Quantification | Sex | UAS control | Gal4 control | Experimental Line | Comparison | Test Used | P Value |
|---|---|---|---|---|---|---|---|---|
| Table 1 (25C) | Table 1. Rhythmic Power | M | *;UAS-NaChBac* | *SLP316 split Gal4 (SS76489)* | *SLP316 > NaChBac* | UAS vs Gal4 | Kruskal-Wallis Test | <0.0001 (adjusted) |
| | | | | | | UAS vs Exp | Kruskal-Wallis Test | <0.0001 (adjusted) |
| | | | | | | Gal4 vs Exp | Kruskal-Wallis Test | >0.9999 (adjusted) |
| 3, Table 1 (25C) | 3B. Free-running period (h) | M | *;UAS-NaChBac* | *SLP316 split Gal4 (SS76489)* | *SLP316 > NaChBac* | UAS vs Gal4 | Kruskal-Wallis Test | 0.1866 (adjusted) |
| | | | | | | UAS vs Exp | Kruskal-Wallis Test | <0.0001 (adjusted) |
| | | | | | | Gal4 vs Exp | Kruskal-Wallis Test | <0.0001 (adjusted) |
| 3, Table 1 (25C) | 3C. Percent Rhythmicity | M | *;UAS-NaChBac* | *SLP316 split Gal4 (SS76489)* | *SLP316 > NaChBac* | UAS vs Gal4 | Fisher's Exact Contingency | >0.2155 (exact) |
| | | | | | | UAS vs Exp | Fisher's Exact Contingency | >0.9999 (exact) |
| | | | | | | Gal4 vs Exp | Fisher's Exact Contingency | 0.2215 (exact) |
| 3 | 3E. DD8 Phase (CT,h) | M | *;UAS-NaChBac* | *SLP316 split Gal4 (SS76489)* | *SLP316 > NaChBac* | UAS vs Gal4 | Kruskal-Wallis Test | 0.5612 (adjusted) |
| | | | | | | UAS vs Exp | Kruskal-Wallis Test | <0.0001 (adjusted) |
| | | | | | | Gal4 vs Exp | Kruskal-Wallis Test | <0.0001 (adjusted) |
| Table 1 (25C) | Table 1. Free-runing period (h) | M | *;pUAS-tetx-LC-TNT;* | *;PdfRed,PdfGal4;* | *PdfRed,PdfGal4; > tetx-LC-TNT* | UAS vs Gal4 | Kruskal-Wallis Test | <0.0001 (adjusted) |
| | | | | | | UAS vs Exp | Kruskal-Wallis Test | <0.0001 (adjusted) |
| | | | | | | Gal4 vs Exp | Kruskal-Wallis Test | <0.0001 (adjusted) |
| 3, Table 1 (25C) | 3E. Percent Rhythmicity | M | *;UAS-NaChBac* | *SLP316 split Gal4 (SS76489)* | *SLP316 > NaChBac* | UAS vs Gal4 | Fisher's Exact Contingency | >0.2155 (exact) |
| | | | | | | UAS vs Exp | Fisher's Exact Contingency | >0.9999 (exact) |
| | | | | | | Gal4 vs Exp | Fisher's Exact Contingency | 0.2215 (exact) |
| 3, Table 1 (25C) | 3G. Free-running period (h) | M | *;UAS-hid;UAS-GFP* | *SLP316 split Gal4 (SS76489)* | *SLP316 > hid+GFP* | UAS vs Gal4 | Kruskal-Wallis Test | >0.9999 (adjusted) |

| | | | | | | | | |
|---|---|---|---|---|---|---|---|---|
| | | | | | | UAS vs Exp | Kruskal-Wallis Test | <0.0001 (adjusted) |
| | | | | | | Gal4 vs Exp | Kruskal-Wallis Test | <0.0001 (adjusted) |
| 3, Table 1 (25C) | 3H. Percent Rhythmicity | M | *;UAS-hid;UAS-GFP* | *SLP316 split Gal4 (SS76489)* | *SLP316 > hid+GFP* | UAS vs Gal4 | Fisher's Exact Contingency | >0.9999 (exact) |
| | | | | | | UAS vs Exp | Fisher's Exact Contingency | 0.4722 (exact) |
| | | | | | | Gal4 vs Exp | Fisher's Exact Contingency | >0.9999 (exact) |
| 3, Table 1 (25C) | 3I. Free-running period (h) | M | *;pUAS-tetx-LC-TNT;* | *SLP316 split Gal4 (SS76489)* | *SLP316 > tetx-LC-TNT* | UAS vs Gal4 | Kruskal-Wallis Test | <0.0001 (adjusted) |
| | | | | | | UAS vs Exp | Kruskal-Wallis Test | <0.0001 (adjusted) |
| | | | | | | Gal4 vs Exp | Kruskal-Wallis Test | 0.0008 (adjusted) |
| 3, Table 1 (25C) | 3J. Percent Rhythmicity | M | *;pUAS-tetx-LC-TNT;* | *SLP316 split Gal4 (SS76489)* | *SLP316 > tetx-LC-TNT* | UAS vs Gal4 | Fisher's Exact Contingency | 0.4435 (exact) |
| | | | | | | UAS vs Exp | Fisher's Exact Contingency | 0.0023 (exact) |
| | | | | | | Gal4 vs Exp | Fisher's Exact Contingency | 0.0002 (exact) |
| 3 | 3L. DD8 Phase (CT,h) | M | *;pUAS-tetx-LC-TNT;* | *SLP316 split Gal4 (SS76489)* | *SLP316 > tetx-LC-TNT* | UAS vs Gal4 | Kruskal-Wallis Test | >0.9999 (exact) |
| | | | | | | UAS vs Exp | Kruskal-Wallis Test | <0.0001 (exact) |
| | | | | | | Gal4 vs Exp | Kruskal-Wallis Test | <0.0001 (exact) |

| **Figure** | **Quantification** | **Sex** | **Control Line** | **Experimental Line** | **Comparison** | **Test Used** | **P Value** |
|---|---|---|---|---|---|---|---|
| 4 | 4B. Eclosion period (h) | N/A | *;UAS-hid;UAS-GFP* | *SLP316 > hid+GFP* | Control vs Experimental | Mann-Whitney Test | 0.9143 (exact) |

| **Figure** | **Quantification** | **Sex** | **UAS control** | **Gal4 control** | **Experimental Line** | **Comparison** | **Test Used** | **P Value** |
|---|---|---|---|---|---|---|---|---|
| 5 | 5B. Daytime activity (counts) | M | *w1118* | *pdf01* | WT x Mutant | Mann-Whitney Test | <0.0001 (exact) | |
| 5 | 5D. Daytime sleep (mins) | M | *w1118* | *pdf01* | WT x Mutant | Mann-Whitney Test | <0.0001 (exact) | |
| 5 | 5F. Daytime activity (counts) | M | *;UAS-hid;UAS-GFP* | *SLP316 split Gal4 (SS76489)* | *SLP316 > hid+GFP* | UAS vs Gal4 | Kruskal-Wallis Test | 0.1586 (adjusted) |
| | | | | | | UAS vs Exp | Kruskal-Wallis Test | <0.0001 (adjusted) |
| | | | | | | Gal4 vs Exp | Kruskal-Wallis Test | <0.0001 (adjusted) |
| 5 | 5H. Daytime sleep (mins) | M | *;UAS-hid;UAS-GFP* | *SLP316 split Gal4 (SS76489)* | *SLP316 > hid+GFP* | UAS vs Gal4 | Kruskal-Wallis Test | 0.0602 (adjusted) |
| | | | | | | UAS vs Exp | Kruskal-Wallis Test | <0.0001 (adjusted) |
| | | | | | | Gal4 vs Exp | Kruskal-Wallis Test | <0.0001 (adjusted) |
| 5 | 5J. Daytime activity (counts) | M | *;pUAS-tetx-LC-TNT;* | *SLP316 split Gal4 (SS76489)* | *SLP316 > tetx-LC-TNT* | UAS vs Gal4 | Kruskal-Wallis Test | 0.0112 (adjusted) |
| | | | | | | UAS vs Exp | Kruskal-Wallis Test | <0.0001 (adjusted) |

| | | | | | | | | |
|---|---|---|---|---|---|---|---|---|
| | | | | | | Gal4 vs Exp | Kruskal-Wallis Test | <0.0001 (adjusted) |
| 5 | 5L. Daytime sleep (mins) | M | *;pUAS-tetx-LC-TNT;* | *SLP316 split Gal4 (SS76489)* | *SLP316 > tetx-LC-TNT* | UAS vs Gal4 | Kruskal-Wallis Test | 0.0107 (adjusted) |
| | | | | | | UAS vs Exp | Kruskal-Wallis Test | <0.0001 (adjusted) |
| | | | | | | Gal4 vs Exp | Kruskal-Wallis Test | <0.0001 (adjusted) |
| **Figure** | **Quantification** | **Sex** | **UAS control** | **Gal4 control** | **Experimental Line** | **Comparison** | **Test Used** | **P Value** |
| 6 | 6B. Daytime sleep (mins) | M | *;UAS-hid;UAS-GFP* | *SLP316 split Gal4 (SS76489)* | *SLP316 > hid+GFP* | UAS vs Gal4 | Kruskal-Wallis Test | <0.0001 (adjusted) |
| | | | | | | UAS vs Exp | Kruskal-Wallis Test | <0.0001 (adjusted) |
| | | | | | | Gal4 vs Exp | Kruskal-Wallis Test | <0.0001 (adjusted) |
| 6 | 6C. P(doze) ZT0-3 | M | *;UAS-hid;UAS-GFP* | *SLP316 split Gal4 (SS76489)* | *SLP316 > hid+GFP* | UAS vs Gal4 | Tukey's Multiple Comparison's | 0.0260 (adjusted) |
| | | | | | | UAS vs Exp | Tukey's Multiple Comparison's | <0.0001 (adjusted) |
| | | | | | | Gal4 vs Exp | Tukey's Multiple Comparison's | 0.0074 (adjusted) |
| 6 | 6C. P(doze) ZT3-6 | M | *;UAS-hid;UAS-GFP* | *SLP316 split Gal4 (SS76489)* | *SLP316 > hid+GFP* | UAS vs Gal4 | Tukey's Multiple Comparison's | 0.0004 (adjusted) |
| | | | | | | UAS vs Exp | Tukey's Multiple Comparison's | <0.0001 (adjusted) |
| | | | | | | Gal4 vs Exp | Tukey's Multiple Comparison's | 0.0361 (adjusted) |
| 6 | 6C. P(doze) ZT6-9 | M | *;UAS-hid;UAS-GFP* | *SLP316 split Gal4 (SS76489)* | *SLP316 > hid+GFP* | UAS vs Gal4 | Tukey's Multiple Comparison's | 0.9501 (adjusted) |
| | | | | | | UAS vs Exp | Tukey's Multiple Comparison's | <0.0001 (adjusted) |
| | | | | | | Gal4 vs Exp | Tukey's Multiple Comparison's | <0.0001 (adjusted) |
| 6 | 6C. P(doze) ZT9-12 | M | *;UAS-hid;UAS-GFP* | *SLP316 split Gal4 (SS76489)* | *SLP316 > hid+GFP* | UAS vs Gal4 | Tukey's Multiple Comparison's | <0.0001 (adjusted) |
| | | | | | | UAS vs Exp | Tukey's Multiple Comparison's | <0.0001 (adjusted) |
| | | | | | | Gal4 vs Exp | Tukey's Multiple Comparison's | 0.0195 (adjusted) |
| 6 | 6C. P(doze) ZT12-15 | M | *;UAS-hid;UAS-GFP* | *SLP316 split Gal4 (SS76489)* | *SLP316 > hid+GFP* | UAS vs Gal4 | Tukey's Multiple Comparison's | <0.0001 (adjusted) |
| | | | | | | UAS vs Exp | Tukey's Multiple Comparison's | <0.0001 (adjusted) |
| | | | | | | Gal4 vs Exp | Tukey's Multiple Comparison's | 0.3970 (adjusted) |
| 6 | 6C. P(doze) ZT15-18 | M | *;UAS-hid;UAS-GFP* | *SLP316 split Gal4 (SS76489)* | *SLP316 > hid+GFP* | UAS vs Gal4 | Tukey's Multiple Comparison's | <0.0001 (adjusted) |
| | | | | | | UAS vs Exp | Tukey's Multiple Comparison's | 0.0078 (adjusted) |
| | | | | | | Gal4 vs Exp | Tukey's Multiple Comparison's | <0.0001 (adjusted) |
| 6 | 6C. P(doze) ZT18-21 | M | *;UAS-hid;UAS-GFP* | *SLP316 split Gal4 (SS76489)* | *SLP316 > hid+GFP* | UAS vs Gal4 | Tukey's Multiple Comparison's | <0.0001 (adjusted) |

| | | | | | | | | |
|---|---|---|---|---|---|---|---|---|
| | | | | | | UAS vs Exp | Tukey's Multiple Comparison's | <0.0001 (adjusted) |
| | | | | | | Gal4 vs Exp | Tukey's Multiple Comparison's | <0.0001 (adjusted) |
| 6 | 6C. P(doze) ZT21-24 | M | *;UAS-hid;UAS-GFP* | *SLP316 split Gal4 (SS76489)* | *SLP316 > hid+GFP* | UAS vs Gal4 | Tukey's Multiple Comparison's | 0.3546 (adjusted) |
| | | | | | | UAS vs Exp | Tukey's Multiple Comparison's | <0.0001 (adjusted) |
| | | | | | | Gal4 vs Exp | Tukey's Multiple Comparison's | <0.0001 (adjusted) |
| 6 | 6D. Average P(doze) | M | *;UAS-hid;UAS-GFP* | *SLP316 split Gal4 (SS76489)* | *SLP316 > hid+GFP* | UAS vs Gal4 | Kruskal-Wallis Test | >0.9999 (adjusted) |
| | | | | | | UAS vs Exp | Kruskal-Wallis Test | <0.0001 (adjusted) |
| | | | | | | Gal4 vs Exp | Kruskal-Wallis Test | <0.0001 (adjusted) |
| 6 | 6F. Daytime sleep (mins) | M | *;pUAS-tetx-LC-TNT;* | *SLP316 split Gal4 (SS76489)* | *SLP316 > tetx-LC-TNT* | UAS vs Gal4 | Kruskal-Wallis Test | >0.9999 (adjusted) |
| | | | | | | UAS vs Exp | Kruskal-Wallis Test | <0.0001 (adjusted) |
| | | | | | | Gal4 vs Exp | Kruskal-Wallis Test | <0.0001 (adjusted) |
| 6 | 6G. P(doze) ZT0-3 | M | *;pUAS-tetx-LC-TNT;* | *SLP316 split Gal4 (SS76489)* | *SLP316 > tetx-LC-TNT* | UAS vs Gal4 | Tukey's Multiple Comparison's | 0.07556 (adjusted) |
| | | | | | | UAS vs Exp | Tukey's Multiple Comparison's | <0.0001 (adjusted) |
| | | | | | | Gal4 vs Exp | Tukey's Multiple Comparison's | <0.0001 (adjusted) |
| 6 | 6G. P(doze) ZT3-6 | M | *;pUAS-tetx-LC-TNT;* | *SLP316 split Gal4 (SS76489)* | *SLP316 > tetx-LC-TNT* | UAS vs Gal4 | Tukey's Multiple Comparison's | <0.0001 (adjusted) |
| | | | | | | UAS vs Exp | Tukey's Multiple Comparison's | <0.0001 (adjusted) |
| | | | | | | Gal4 vs Exp | Tukey's Multiple Comparison's | 0.7333 (adjusted) |
| 6 | 6G. P(doze) ZT6-9 | M | *;pUAS-tetx-LC-TNT;* | *SLP316 split Gal4 (SS76489)* | *SLP316 > tetx-LC-TNT* | UAS vs Gal4 | Tukey's Multiple Comparison's | 0.0851 (adjusted) |
| | | | | | | UAS vs Exp | Tukey's Multiple Comparison's | <0.0001 (adjusted) |
| | | | | | | Gal4 vs Exp | Tukey's Multiple Comparison's | 0.0002 (adjusted) |
| 6 | 6G. P(doze) ZT9-12 | M | *;pUAS-tetx-LC-TNT;* | *SLP316 split Gal4 (SS76489)* | *SLP316 > tetx-LC-TNT* | UAS vs Gal4 | Tukey's Multiple Comparison's | 0.0176 (adjusted) |
| | | | | | | UAS vs Exp | Tukey's Multiple Comparison's | <0.0001 (adjusted) |
| | | | | | | Gal4 vs Exp | Tukey's Multiple Comparison's | <0.0001 (adjusted) |
| 6 | 6G. P(doze) ZT12-15 | M | *;pUAS-tetx-LC-TNT;* | *SLP316 split Gal4 (SS76489)* | *SLP316 > tetx-LC-TNT* | UAS vs Gal4 | Tukey's Multiple Comparison's | 0.0652 (adjusted) |
| | | | | | | UAS vs Exp | Tukey's Multiple Comparison's | <0.0001 (adjusted) |
| | | | | | | Gal4 vs Exp | Tukey's Multiple Comparison's | <0.0001 (adjusted) |
| 6 | 6G. P(doze) ZT15-18 | M | *;pUAS-tetx-LC-TNT;* | *SLP316 split Gal4 (SS76489)* | *SLP316 > tetx-LC-TNT* | UAS vs Gal4 | Tukey's Multiple Comparison's | <0.0001 (adjusted) |

| | | | | | | | | |
|---|---|---|---|---|---|---|---|---|
| | | | | | | UAS vs Exp | Tukey's Multiple Comparison's | 0.0749 (adjusted) |
| | | | | | | Gal4 vs Exp | Tukey's Multiple Comparison's | <0.0001 (adjusted) |
| 6 | 6G. P(doze) ZT18-21 | M | *;pUAS-tetx-LC-TNT;* | *SLP316 split Gal4 (SS76489)* | *SLP316 > tetx-LC-TNT* | UAS vs Gal4 | Tukey's Multiple Comparison's | <0.0001 (adjusted) |
| | | | | | | UAS vs Exp | Tukey's Multiple Comparison's | 0.0002 (adjusted) |
| | | | | | | Gal4 vs Exp | Tukey's Multiple Comparison's | <0.0001 (adjusted) |
| 6 | 6G. P(doze) ZT21-24 | M | *;pUAS-tetx-LC-TNT;* | *SLP316 split Gal4 (SS76489)* | *SLP316 > tetx-LC-TNT* | UAS vs Gal4 | Tukey's Multiple Comparison's | 0.9988 (adjusted) |
| | | | | | | UAS vs Exp | Tukey's Multiple Comparison's | <0.0001 (adjusted) |
| | | | | | | Gal4 vs Exp | Tukey's Multiple Comparison's | <0.0001 (adjusted) |
| 6 | 6H. Average P(doze) | M | *;pUAS-tetx-LC-TNT;* | *SLP316 split Gal4 (SS76489)* | *SLP316 > tetx-LC-TNT* | UAS vs Gal4 | Kruskal-Wallis Test | >0.9999 (adjusted) |
| | | | | | | UAS vs Exp | Kruskal-Wallis Test | <0.0001 (adjusted) |
| | | | | | | Gal4 vs Exp | Kruskal-Wallis Test | <0.0001 (adjusted) |
| Table 1 (25C) | Table 1. Percent Rhythmicity | M | *;pUAS-tetx-LC-TNT;* | *;PdfRed,PdfGal4;* | *PdfRed,PdfGal4; > tetx-LC-TNT* | UAS vs Gal4 | Fisher's Exact Contingency | 0.1063 (exact) |
| | | | | | | UAS vs Exp | Fisher's Exact Contingency | 0.0010 (exact) |
| | | | | | | Gal4 vs Exp | Fisher's Exact Contingency | <0.0001 (exact) |
| Table 1 (25C) | Table 1. Rhythmic Power | M | *;pUAS-tetx-LC-TNT;* | *;PdfRed,PdfGal4;* | *PdfRed,PdfGal4; > tetx-LC-TNT* | UAS vs Gal4 | Kruskal-Wallis Test | <0.0001 (adjusted) |
| | | | | | | UAS vs Exp | Kruskal-Wallis Test | 0.0219 (adjusted) |
| | | | | | | Gal4 vs Exp | Kruskal-Wallis Test | <0.0001 (adjusted) |
| Table 1 (25C) | Table 1. Rhythmic Power | M | *;UAS-hid;UAS-GFP* | *SLP316 split Gal4 (SS76489)* | *SLP316 > hid+GFP* | UAS vs Gal4 | Kruskal-Wallis Test | <0.0001 (adjusted) |
| | | | | | | UAS vs Exp | Kruskal-Wallis Test | 0.0319 (adjusted) |
| | | | | | | Gal4 vs Exp | Kruskal-Wallis Test | <0.0001 (adjusted) |
| Table 1 (25C) | Table 1. Rhythmic Power | M | *;pUAS-tetx-LC-TNT;* | *SLP316 split Gal4 (SS76489)* | *SLP316 > tetx-LC-TNT* | UAS vs Gal4 | Kruskal-Wallis Test | 0.0287 (adjusted) |
| | | | | | | UAS vs Exp | Kruskal-Wallis Test | <0.0001 (adjusted) |
| | | | | | | Gal4 vs Exp | Kruskal-Wallis Test | <0.0001 (adjusted) |
| Table 1 (25C) | Table 1. Rhythmic Power | F | *;UAS-hid;UAS-GFP* | *SLP316 split Gal4 (SS76489)* | *SLP316 > hid+GFP* | UAS vs Gal4 | Kruskal-Wallis Test | <0.0001 (adjusted) |
| | | | | | | UAS vs Exp | Kruskal-Wallis Test | >0.9999 (adjusted) |
| | | | | | | Gal4 vs Exp | Kruskal-Wallis Test | <0.0001 (adjusted) |
| Table 1 (25C) | Table 1. Rhythmic Power | F | *;pUAS-tetx-LC-TNT;* | *SLP316 split Gal4 (SS76489)* | *SLP316 > tetx-LC-TNT* | UAS vs Gal4 | Kruskal-Wallis Test | 0.3084 (adjusted) |

| | | | | | | | | |
|---|---|---|---|---|---|---|---|---|
| | | | | | | UAS vs Exp | Kruskal-Wallis Test | >0.9999 (adjusted) |
| | | | | | | Gal4 vs Exp | Kruskal-Wallis Test | 0.0318 (adjusted) |
| Table 1 (18C) | Table 1. Percent Rhythmicity | M | *;UAS-NaChBac* | *SLP316 split Gal4 (SS76489)* | *SLP316 > NaChBac* | UAS vs Gal4 | Fisher's Exact Contingency | >0.9999 (exact) |
| | | | | | | UAS vs Exp | Fisher's Exact Contingency | >0.9999 (exact) |
| | | | | | | Gal4 vs Exp | Fisher's Exact Contingency | >0.9999 (exact) |
| Table 1 (18C) | Table 1. Free-running period (h) | M | *;UAS-NaChBac* | *SLP316 split Gal4 (SS76489)* | *SLP316 > NaChBac* | UAS vs Gal4 | Kruskal-Wallis Test | 0.0010 (adjusted) |
| | | | | | | UAS vs Exp | Kruskal-Wallis Test | 0.7869 (adjusted) |
| | | | | | | Gal4 vs Exp | Kruskal-Wallis Test | 0.3485 (adjusted) |
| Table 1 (18C) | Table 1. Rhythmic Power | M | *;UAS-NaChBac* | *SLP316 split Gal4 (SS76489)* | *SLP316 > NaChBac* | UAS vs Gal4 | Kruskal-Wallis Test | >0.9999 (adjusted) |
| | | | | | | UAS vs Exp | Kruskal-Wallis Test | >0.9999 (adjusted) |
| | | | | | | Gal4 vs Exp | Kruskal-Wallis Test | >0.9999 (adjusted) |
| Table 1 (18C) | Table 1. Percent Rhythmicity | M | *;UAS-hid;UAS-GFP* | *SLP316 split Gal4 (SS76489)* | *SLP316 > hid+GFP* | UAS vs Gal4 | Fisher's Exact Contingency | 0.2461 (exact) |
| | | | | | | UAS vs Exp | Fisher's Exact Contingency | 0.0007 (exact) |
| | | | | | | Gal4 vs Exp | Fisher's Exact Contingency | 0.0453 (exact) |
| Table 1 (18C) | Table 1. Free-running period (h) | M | *;UAS-hid;UAS-GFP* | *SLP316 split Gal4 (SS76489)* | *SLP316 > hid+GFP* | UAS vs Gal4 | Kruskal-Wallis Test | 0.3144 (adjusted) |
| | | | | | | UAS vs Exp | Kruskal-Wallis Test | <0.0001 (adjusted) |
| | | | | | | Gal4 vs Exp | Kruskal-Wallis Test | <0.0001 (adjusted) |
| Table 1 (18C) | Table 1. Rhythmic Power | M | *;UAS-hid;UAS-GFP* | *SLP316 split Gal4 (SS76489)* | *SLP316 > hid+GFP* | UAS vs Gal4 | Kruskal-Wallis Test | 0.7079 (adjusted) |
| | | | | | | UAS vs Exp | Kruskal-Wallis Test | <0.0001 (adjusted) |
| | | | | | | Gal4 vs Exp | Kruskal-Wallis Test | 0.2999 (adjusted) |
| Table 1 (18C) | Table 1. Percent Rhythmicity | M | *;pUAS-tetx-LC-TNT;* | *SLP316 split Gal4 (SS76489)* | *SLP316 > tetx-LC-TNT* | UAS vs Gal4 | Fisher's Exact Contingency | 0.0007 (exact) |
| | | | | | | UAS vs Exp | Fisher's Exact Contingency | 0.0967 (exact) |
| | | | | | | Gal4 vs Exp | Fisher's Exact Contingency | <0.0001 (exact) |
| Table 1 (18C) | Table 1. Free-running period (h) | M | *;pUAS-tetx-LC-TNT;* | *SLP316 split Gal4 (SS76489)* | *SLP316 > tetx-LC-TNT* | UAS vs Gal4 | Kruskal-Wallis Test | 0.9943 (adjusted) |
| | | | | | | UAS vs Exp | Kruskal-Wallis Test | <0.0001 (adjusted) |
| | | | | | | Gal4 vs Exp | Kruskal-Wallis Test | <0.0001 (adjusted) |
| Table 1 (18C) | Table 1. Rhythmic Power | M | *;pUAS-tetx-LC-TNT;* | *SLP316 split Gal4 (SS76489)* | *SLP316 > tetx-LC-TNT* | UAS vs Gal4 | Kruskal-Wallis Test | <0.0001 (adjusted) |

| | | | | | | | | |
|---|---|---|---|---|---|---|---|---|
| | | | | | | UAS vs Exp | Kruskal-Wallis Test | 0.0468 (adjusted) |
| | | | | | | Gal4 vs Exp | Kruskal-Wallis Test | <0.0001 (adjusted) |
| **Figure** | **Quantification** | **Sex** | **Condition 1** | **Condition 2** | **Comparison** | **Test Used** | **P Value** | |
| S2 | S2B. Cell body count | M | 1 week old | 5 week old | 1 week vs 5 weeks old | Mann-Whitney Test | 0.3053 | |
| **Figure** | **Quantification** | **Sex** | **UAS control** | **Gal4 control** | **Experimental Line** | **Comparison** | **Test Used** | **P Value** |
| S3 | S4. Daytime activity (counts) | M | *;UAS-NaChBac* | *SLP316 split Gal4 (SS76489)* | *SLP316 > NaChBac* | UAS vs Gal4 | Kruskal-Wallis Test | 0.0119 (adjusted) |
| | | | | | | UAS vs Exp | Kruskal-Wallis Test | <0.0001 (adjusted) |
| | | | | | | Gal4 vs Exp | Kruskal-Wallis Test | <0.0001 (adjusted) |
| S3 | S3C. Morning peak time (ZT, h) | M | *;UAS-NaChBac* | *SLP316 split Gal4 (SS76489)* | *SLP316 > NaChBac* | UAS vs Gal4 | Kruskal-Wallis Test | <0.0001 (adjusted) |
| | | | | | | UAS vs Exp | Kruskal-Wallis Test | <0.0001 (adjusted) |
| | | | | | | Gal4 vs Exp | Kruskal-Wallis Test | 0.9878 (adjusted) |
| S3 | S3D. Evening peak time (ZT, h) | M | *;UAS-NaChBac* | *SLP316 split Gal4 (SS76489)* | *SLP316 > NaChBac* | UAS vs Gal4 | Kruskal-Wallis Test | 0.8415 (adjusted) |
| | | | | | | UAS vs Exp | Kruskal-Wallis Test | 0.0269 (adjusted) |
| | | | | | | Gal4 vs Exp | Kruskal-Wallis Test | 0.3266 (adjusted) |
| S3 | S3E. Evening peak area | M | *;UAS-NaChBac* | *SLP316 split Gal4 (SS76489)* | *SLP316 > NaChBac* | UAS vs Gal4 | Kruskal-Wallis Test | <0.0001 (adjusted) |
| | | | | | | UAS vs Exp | Kruskal-Wallis Test | 0.1582 (adjusted) |
| | | | | | | Gal4 vs Exp | Kruskal-Wallis Test | <0.0001 (adjusted) |
| S3 | S3G. Daytime activity (counts) | M | *;UAS-hid;UAS-GFP* | *SLP316 split Gal4 (SS76489)* | *SLP316 > hid+GFP* | UAS vs Gal4 | Kruskal-Wallis Test | 0.2165 (adjusted) |
| | | | | | | UAS vs Exp | Kruskal-Wallis Test | <0.0001 (adjusted) |
| | | | | | | Gal4 vs Exp | Kruskal-Wallis Test | <0.0001 (adjusted) |
| S3 | S3H. Morning peak time (ZT, h) | M | *;UAS-hid;UAS-GFP* | *SLP316 split Gal4 (SS76489)* | *SLP316 > hid+GFP* | UAS vs Gal4 | Kruskal-Wallis Test | <0.0001 (adjusted) |
| | | | | | | UAS vs Exp | Kruskal-Wallis Test | <0.0001 (adjusted) |
| | | | | | | Gal4 vs Exp | Kruskal-Wallis Test | 0.0001 (adjusted) |
| S3 | S3I. Evening peak time (ZT, h) | M | *;UAS-hid;UAS-GFP* | *SLP316 split Gal4 (SS76489)* | *SLP316 > hid+GFP* | UAS vs Gal4 | Kruskal-Wallis Test | <0.0001 (adjusted) |
| | | | | | | UAS vs Exp | Kruskal-Wallis Test | <0.0001 (adjusted) |
| | | | | | | Gal4 vs Exp | Kruskal-Wallis Test | 0.4879 (adjusted) |
| S3 | S3J. Evening peak area | M | *;UAS-hid;UAS-GFP* | *SLP316 split Gal4 (SS76489)* | *SLP316 > hid+GFP* | UAS vs Gal4 | Kruskal-Wallis Test | <0.0001 (adjusted) |

| | | | | | | | | |
|---|---|---|---|---|---|---|---|---|
| | | | | | | UAS vs Exp | Kruskal-Wallis Test | <0.0001 (adjusted) |
| | | | | | | Gal4 vs Exp | Kruskal-Wallis Test | <0.0001 (adjusted) |
| S3 | S3L. Daytime activity (counts) | M | *;pUAS-tetx-LC-TNT;* | *SLP316 split Gal4 (SS76489)* | *SLP316 > tetx-LC-TNT* | UAS vs Gal4 | Kruskal-Wallis Test | <0.0001 (adjusted) |
| | | | | | | UAS vs Exp | Kruskal-Wallis Test | <0.0001 (adjusted) |
| | | | | | | Gal4 vs Exp | Kruskal-Wallis Test | <0.0001 (adjusted) |
| S3 | S3M. Morning peak time (ZT, h) | M | *;pUAS-tetx-LC-TNT;* | *SLP316 split Gal4 (SS76489)* | *SLP316 > tetx-LC-TNT* | UAS vs Gal4 | Kruskal-Wallis Test | <0.0001 (adjusted) |
| | | | | | | UAS vs Exp | Kruskal-Wallis Test | 0.0077 (adjusted) |
| | | | | | | Gal4 vs Exp | Kruskal-Wallis Test | 0.0001 (adjusted) |
| S3 | S3N. Evening peak time (ZT, h) | M | *;pUAS-tetx-LC-TNT;* | *SLP316 split Gal4 (SS76489)* | *SLP316 > tetx-LC-TNT* | UAS vs Gal4 | Kruskal-Wallis Test | 0.0005 (adjusted) |
| | | | | | | UAS vs Exp | Kruskal-Wallis Test | 0.3420 (adjusted) |
| | | | | | | Gal4 vs Exp | Kruskal-Wallis Test | 0.0898 (adjusted) |
| S3 | S3O. Evening peak area | M | *;pUAS-tetx-LC-TNT;* | *SLP316 split Gal4 (SS76489)* | *SLP316 > tetx-LC-TNT* | UAS vs Gal4 | Kruskal-Wallis Test | >0.9999 (adjusted) |
| | | | | | | UAS vs Exp | Kruskal-Wallis Test | <0.0001 (adjusted) |
| | | | | | | Gal4 vs Exp | Kruskal-Wallis Test | <0.0001 (adjusted) |
| S3 | S3Q. DD8 Phase (CT, h) | M | *;UAS-hid;UAS-GFP* | *SLP316 split Gal4 (SS76489)* | *SLP316 > hid+GFP* | UAS vs Gal4 | Kruskal-Wallis Test | <0.0001 (adjusted) |
| | | | | | | UAS vs Exp | Kruskal-Wallis Test | 0.0261 (adjusted) |
| | | | | | | Gal4 vs Exp | Kruskal-Wallis Test | <0.0001 (adjusted) |
| **Figure** | **Quantification** | **Sex** | **UAS control** | **Gal4 control** | **Experimental Line** | **Test Used** | **P Value** | |
| S4, Table 1 (25C) | S4A. Free-running period (h) | F | *;UAS-hid;UAS-GFP* | *SLP316 split Gal4 (SS76489)* | *SLP316 > hid+GFP* | UAS vs Gal4 | Kruskal-Wallis Test | >0.9999 (adjusted) |
| | | | | | | UAS vs Exp | Kruskal-Wallis Test | 0.0004 (adjusted) |
| | | | | | | Gal4 vs Exp | Kruskal-Wallis Test | 0.0154 (adjusted) |
| S4, Table 1 (25C) | S4C. Free-running period (h) | F | *;pUAS-tetx-LC-TNT;* | *SLP316 split Gal4 (SS76489)* | *SLP316 > tetx-LC-TNT* | UAS vs Gal4 | Kruskal-Wallis Test | 0.0005 (adjusted) |
| | | | | | | UAS vs Exp | Kruskal-Wallis Test | <0.0001 (adjusted) |
| | | | | | | Gal4 vs Exp | Kruskal-Wallis Test | >0.9999 (adjusted) |
| S4 | S4B. Difference in FRP | F vs M | *SLP316 > hid+GFP* | *SLP316 > hid+GFP* | Female x Male | Mann-Whitney Test | 0.6673 | |
| S4 | S4D. Difference in FRP | F vs M | *SLP316 > tetx-LC-TNT* | *SLP316 > tetx-LC-TNT* | Female x Male | Mann-Whitney Test | 0.1971 | |
| S4, Table 1 (25C) | S4E. Percent Rhythmicity | F | *;UAS-hid;UAS-GFP* | *SLP316 split Gal4 (SS76489)* | *SLP316 > hid+GFP* | UAS vs Gal4 | Fisher's Exact Contingency | 0.0108 (exact) |

| | | | | | | | | |
|---|---|---|---|---|---|---|---|---|
| | | | | | | UAS vs Exp | Fisher's Exact Contingency | >0.9999 (exact) |
| | | | | | | Gal4 vs Exp | Fisher's Exact Contingency | 0.0108 (exact) |
| S4, Table 1 (25C) | S4E. Percent Rhythmicity | F | *;pUAS-tetx-LC-TNT;* | *SLP316 split Gal4 (SS76489)* | *SLP316 > tetx-LC-TNT* | UAS vs Gal4 | Fisher's Exact Contingency | 0.2566 (exact) |
| | | | | | | UAS vs Exp | Fisher's Exact Contingency | 0.6059 (exact) |
| | | | | | | Gal4 vs Exp | Fisher's Exact Contingency | 0.1035 (exact) |
| S4 | S4G. Daytime sleep (mins) | F | *;UAS-hid;UAS-GFP* | *SLP316 split Gal4 (SS76489)* | *SLP316 > hid+GFP* | UAS vs Gal4 | Kruskal-Wallis Test | 0.2822 (adjusted) |
| | | | | | | UAS vs Exp | Kruskal-Wallis Test | <0.0001 (adjusted) |
| | | | | | | Gal4 vs Exp | Kruskal-Wallis Test | 0.0002 (adjusted) |
| S4 | S4H. Nighttime sleep (mins) | F | *;UAS-hid;UAS-GFP* | *SLP316 split Gal4 (SS76489)* | *SLP316 > hid+GFP* | UAS vs Gal4 | Kruskal-Wallis Test | <0.0001 (adjusted) |
| | | | | | | UAS vs Exp | Kruskal-Wallis Test | 0.3044 (adjusted) |
| | | | | | | Gal4 vs Exp | Kruskal-Wallis Test | <0.0001 (adjusted) |
| S4 | S4J. Daytime sleep (mins) | F | *;pUAS-tetx-LC-TNT;* | *SLP316 split Gal4 (SS76489)* | *SLP316 > tetx-LC-TNT* | UAS vs Gal4 | Kruskal-Wallis Test | 0.7029 (adjusted) |
| | | | | | | UAS vs Exp | Kruskal-Wallis Test | 0.0473 (adjusted) |
| | | | | | | Gal4 vs Exp | Kruskal-Wallis Test | 0.0010 (adjusted) |
| S4 | S4K. Nighttime sleep (mins) | F | *;pUAS-tetx-LC-TNT;* | *SLP316 split Gal4 (SS76489)* | *SLP316 > tetx-LC-TNT* | UAS vs Gal4 | Kruskal-Wallis Test | 0.1559 (adjusted) |
| | | | | | | UAS vs Exp | Kruskal-Wallis Test | 0.0003 (adjusted) |
| | | | | | | Gal4 vs Exp | Kruskal-Wallis Test | <0.0001 (adjusted) |
| **Figure** | **Quantification** | **Sex** | **UAS control** | **Gal4 control** | **Experimental Line** | **Test Used** | **P Value** | |
| S5 | S4B. Eclosion period (h) | N/A | *UAS-rpr* | *SLP316 split Gal4 (SS76489)* | *SLP316 > rpr* | UAS vs Gal4 | Kruskal-Wallis Test | 0.5324 (adjusted) |
| | | | | | | UAS vs Exp | Kruskal-Wallis Test | 0.9109 (adjusted) |
| | | | | | | Gal4 vs Exp | Kruskal-Wallis Test | 0.7190 (adjusted) |
| **Figure** | **Quantification** | **Sex** | **UAS control** | **Gal4 control** | **Experimental Line** | **Test Used** | **P Value** | |
| S6 | S6A. Nighttime activity (counts) | M | *;UAS-hid;UAS-GFP* | *SLP316 split Gal4 (SS76489)* | *SLP316 > hid+GFP* | UAS vs Gal4 | Kruskal-Wallis Test | <0.0001 (adjusted) |
| | | | | | | UAS vs Exp | Kruskal-Wallis Test | 0.1512 (adjusted) |
| | | | | | | Gal4 vs Exp | Kruskal-Wallis Test | <0.0001 (adjusted) |
| S6 | S6B. Nighttime sleep (mins) | M | *;UAS-hid;UAS-GFP* | *SLP316 split Gal4 (SS76489)* | *SLP316 > hid+GFP* | UAS vs Gal4 | Kruskal-Wallis Test | <0.0001 (adjusted) |
| | | | | | | UAS vs Exp | Kruskal-Wallis Test | 0.0086 (adjusted) |

| | | | | | | | | |
|---|---|---|---|---|---|---|---|---|
| | | | | | | Gal4 vs Exp | Kruskal-Wallis Test | <0.0001 (adjusted) |
| S6 | S6C. Nighttime activity (counts) | M | *;pUAS-tetx-LC-TNT;* | *SLP316 split Gal4 (SS76489)* | *SLP316 > tetx-LC-TNT* | UAS vs Gal4 | Kruskal-Wallis Test | <0.0001 (adjusted) |
| | | | | | | UAS vs Exp | Kruskal-Wallis Test | 0.8491 (adjusted) |
| | | | | | | Gal4 vs Exp | Kruskal-Wallis Test | <0.0001 (adjusted) |
| S6 | S6D. Nighttime sleep (mins) | M | *;pUAS-tetx-LC-TNT;* | *SLP316 split Gal4 (SS76489)* | *SLP316 > tetx-LC-TNT* | UAS vs Gal4 | Kruskal-Wallis Test | <0.0001 (adjusted) |
| | | | | | | UAS vs Exp | Kruskal-Wallis Test | 0.3436 (adjusted) |
| | | | | | | Gal4 vs Exp | Kruskal-Wallis Test | <0.0001 (adjusted) |

| **Figure** | **Quantification** | **Sex** | **Control Line** | **Experimental Line** | **Comparison** | **Test Used** | **P Value** |
|---|---|---|---|---|---|---|---|
| S6 | S6E. Nihgttime activity (counts) | M | *w1118* | *pdf01* | WT x Mutant | Mann-Whitney Test | 0.0229 (exact) |
| S6 | S6F. Nighttime sleep (mins) | M | *w1118* | *pdf01* | WT x Mutant | Mann-Whitney Test | 0.2013 (exact) |

| **Figure** | **Quantification** | **Sex** | **UAS control** | **Gal4 control** | **Experimental Line** | **Comparison** | **Test Used** | **P Value** |
|---|---|---|---|---|---|---|---|---|
| S7 | S7A. P(wake) ZT0-3 | M | *;UAS-hid;UAS-GFP* | *SLP316 split Gal4 (SS76489)* | *SLP316 > hid+GFP* | UAS vs Gal4 | Tukey's Multiple Comparison's | 0.7904 (adjusted) |
| | | | | | | UAS vs Exp | Tukey's Multiple Comparison's | 0.4127 (adjusted) |
| | | | | | | Gal4 vs Exp | Tukey's Multiple Comparison's | 0.1313 (adjusted) |
| S7 | S7A. P(wake) ZT3-S7 | M | *;UAS-hid;UAS-GFP* | *SLP316 split Gal4 (SS76489)* | *SLP316 > hid+GFP* | UAS vs Gal4 | Tukey's Multiple Comparison's | 0.4429 (adjusted) |
| | | | | | | UAS vs Exp | Tukey's Multiple Comparison's | 0.5848 (adjusted) |
| | | | | | | Gal4 vs Exp | Tukey's Multiple Comparison's | 0.0750 (adjusted) |
| S7 | S7A. P(wake) ZT6-9 | M | *;UAS-hid;UAS-GFP* | *SLP316 split Gal4 (SS76489)* | *SLP316 > hid+GFP* | UAS vs Gal4 | Tukey's Multiple Comparison's | <0.0001 (adjusted) |
| | | | | | | UAS vs Exp | Tukey's Multiple Comparison's | 0.7292 (adjusted) |
| | | | | | | Gal4 vs Exp | Tukey's Multiple Comparison's | <0.0001 (adjusted) |
| S7 | S7A. P(wake) ZT9-12 | M | *;UAS-hid;UAS-GFP* | *SLP316 split Gal4 (SS76489)* | *SLP316 > hid+GFP* | UAS vs Gal4 | Tukey's Multiple Comparison's | <0.0001 (adjusted) |
| | | | | | | UAS vs Exp | Tukey's Multiple Comparison's | <0.0001 (adjusted) |
| | | | | | | Gal4 vs Exp | Tukey's Multiple Comparison's | <0.0001 (adjusted) |
| S7 | S7A. P(wake) ZT12-15 | M | *;UAS-hid;UAS-GFP* | *SLP316 split Gal4 (SS76489)* | *SLP316 > hid+GFP* | UAS vs Gal4 | Tukey's Multiple Comparison's | <0.0001 (adjusted) |
| | | | | | | UAS vs Exp | Tukey's Multiple Comparison's | <0.0001 (adjusted) |
| | | | | | | Gal4 vs Exp | Tukey's Multiple Comparison's | <0.0001 (adjusted) |
| S7 | S7A. P(wake) ZT15-18 | M | *;UAS-hid;UAS-GFP* | *SLP316 split Gal4 (SS76489)* | *SLP316 > hid+GFP* | UAS vs Gal4 | Tukey's Multiple Comparison's | <0.0001 (adjusted) |

| | | | | | | | | |
|---|---|---|---|---|---|---|---|---|
| | | | | | | UAS vs Exp | Tukey's Multiple Comparison's | 0.4129 (adjusted) |
| | | | | | | Gal4 vs Exp | Tukey's Multiple Comparison's | <0.0001 (adjusted) |
| S7 | S7A. P(wake) ZT18-21 | M | *;UAS-hid;UAS-GFP* | *SLP316 split Gal4 (SS76489)* | *SLP316 > hid+GFP* | UAS vs Gal4 | Tukey's Multiple Comparison's | <0.0001 (adjusted) |
| | | | | | | UAS vs Exp | Tukey's Multiple Comparison's | 0.2568 (adjusted) |
| | | | | | | Gal4 vs Exp | Tukey's Multiple Comparison's | <0.0001 (adjusted) |
| S7 | S7A. P(wake) ZT21-24 | M | *;UAS-hid;UAS-GFP* | *SLP316 split Gal4 (SS76489)* | *SLP316 > hid+GFP* | UAS vs Gal4 | Tukey's Multiple Comparison's | <0.0001 (adjusted) |
| | | | | | | UAS vs Exp | Tukey's Multiple Comparison's | <0.0001 (adjusted) |
| | | | | | | Gal4 vs Exp | Tukey's Multiple Comparison's | <0.0001 (adjusted) |
| S7 | S7B. Average P(wake) | M | *;UAS-hid;UAS-GFP* | *SLP316 split Gal4 (SS76489)* | *SLP316 > hid+GFP* | UAS vs Gal4 | Kruskal-Wallis Test | <0.0001 (adjusted) |
| | | | | | | UAS vs Exp | Kruskal-Wallis Test | 0.6120 (adjusted) |
| | | | | | | Gal4 vs Exp | Kruskal-Wallis Test | <0.0001 (adjusted) |
| S7 | S7C. P(wake) ZT0-3 | M | *;pUAS-tetx-LC-TNT;* | *SLP316 split Gal4 (SS76489)* | *SLP316 > tetx-LC-TNT* | UAS vs Gal4 | Tukey's Multiple Comparison's | 0.0502 (adjusted) |
| | | | | | | UAS vs Exp | Tukey's Multiple Comparison's | <0.0001 (adjusted) |
| | | | | | | Gal4 vs Exp | Tukey's Multiple Comparison's | 0.0428 (adjusted) |
| S7 | S7C. P(wake) ZT3-6 | M | *;pUAS-tetx-LC-TNT;* | *SLP316 split Gal4 (SS76489)* | *SLP316 > tetx-LC-TNT* | UAS vs Gal4 | Tukey's Multiple Comparison's | 0.0019 (adjusted) |
| | | | | | | UAS vs Exp | Tukey's Multiple Comparison's | <0.0001 (adjusted) |
| | | | | | | Gal4 vs Exp | Tukey's Multiple Comparison's | 0.1666 (adjusted) |
| S7 | S7C. P(wake) ZT6-9 | M | *;pUAS-tetx-LC-TNT;* | *SLP316 split Gal4 (SS76489)* | *SLP316 > tetx-LC-TNT* | UAS vs Gal4 | Tukey's Multiple Comparison's | 0.0980 (adjusted) |
| | | | | | | UAS vs Exp | Tukey's Multiple Comparison's | 0.6173 (adjusted) |
| | | | | | | Gal4 vs Exp | Tukey's Multiple Comparison's | 0.0074 (adjusted) |
| S7 | S7C. P(wake) ZT9-12 | M | *;pUAS-tetx-LC-TNT;* | *SLP316 split Gal4 (SS76489)* | *SLP316 > tetx-LC-TNT* | UAS vs Gal4 | Tukey's Multiple Comparison's | <0.0001 (adjusted) |
| | | | | | | UAS vs Exp | Tukey's Multiple Comparison's | 0.0922 (adjusted) |
| | | | | | | Gal4 vs Exp | Tukey's Multiple Comparison's | <0.0001 (adjusted) |
| S7 | S7C. P(wake) ZT12-15 | M | *;pUAS-tetx-LC-TNT;* | *SLP316 split Gal4 (SS76489)* | *SLP316 > tetx-LC-TNT* | UAS vs Gal4 | Tukey's Multiple Comparison's | <0.0001 (adjusted) |
| | | | | | | UAS vs Exp | Tukey's Multiple Comparison's | 0.0002 (adjusted) |
| | | | | | | Gal4 vs Exp | Tukey's Multiple Comparison's | <0.0001 (adjusted) |
| S7 | S7C. P(wake) ZT15-18 | M | *;pUAS-tetx-LC-TNT;* | *SLP316 split Gal4 (SS76489)* | *SLP316 > tetx-LC-TNT* | UAS vs Gal4 | Tukey's Multiple Comparison's | <0.0001 (adjusted) |

| | | | | | | | | |
|---|---|---|---|---|---|---|---|---|
| | | | | | | UAS vs Exp | Tukey's Multiple Comparison's | 0.2984 (adjusted) |
| | | | | | | Gal4 vs Exp | Tukey's Multiple Comparison's | <0.0001 (adjusted) |
| S7 | S7C. P(wake) ZT18-21 | M | *;pUAS-tetx-LC-TNT;* | *SLP316 split Gal4 (SS76489)* | *SLP316 > tetx-LC-TNT* | UAS vs Gal4 | Tukey's Multiple Comparison's | <0.0001 (adjusted) |
| | | | | | | UAS vs Exp | Tukey's Multiple Comparison's | 0.4608 (adjusted) |
| | | | | | | Gal4 vs Exp | Tukey's Multiple Comparison's | <0.0001 (adjusted) |
| S7 | S7C. P(wake) ZT21-24 | M | *;pUAS-tetx-LC-TNT;* | *SLP316 split Gal4 (SS76489)* | *SLP316 > tetx-LC-TNT* | UAS vs Gal4 | Tukey's Multiple Comparison's | <0.0001 (adjusted) |
| | | | | | | UAS vs Exp | Tukey's Multiple Comparison's | 0.0227 (adjusted) |
| | | | | | | Gal4 vs Exp | Tukey's Multiple Comparison's | <0.0001 (adjusted) |
| S7 | S7D. Average P(wake) | M | *;pUAS-tetx-LC-TNT;* | *SLP316 split Gal4 (SS76489)* | *SLP316 > tetx-LC-TNT* | UAS vs Gal4 | Kruskal-Wallis Test | <0.0001 (adjusted) |
| | | | | | | UAS vs Exp | Kruskal-Wallis Test | >0.9999 (adjusted) |
| | | | | | | Gal4 vs Exp | Kruskal-Wallis Test | <0.0001 (adjusted) |
